# Prenylation controls proliferation in human pluripotent stem cell-derived cardiomyocytes

**DOI:** 10.1101/2024.07.01.601625

**Authors:** Christopher A.P. Batho, Janice D. Reid, Harley R. Robinson, Henrietta Cserne Szappanos, Lynn A.C. Devilée, Sharon M. Hoyte, Rebecca L. Johnston, Rebekah Ziegman, Sarah Hassan, Lior Soday, Rebecca L. Fitzsimmons, Simon R. Foster, Dominic C. H. Ng, Edward Tate, Enzo R. Porrello, Benjamin L. Parker, Richard J. Mills, James E. Hudson

## Abstract

Induction of cardiomyocyte proliferation to replace damaged heart tissue is a promising therapeutic approach. A recent drug screen revealed that cardiomyocytes require the mevalonate pathway for proliferation, although the specific mechanisms are unknown. In this study, we use human pluripotent stem cell-derived cardiomyocytes and cardiac organoids to further interrogate the role of the mevalonate pathway in cardiomyocyte proliferation. Chemical and genetic perturbations of the mevalonate pathway indicated that the post-translational modification, prenylation, regulates cardiomyocyte proliferation. We use prenyl probes and mass spectrometry to identify a catalogue of 40 prenylated proteins in human cardiac cells, including proteins where prenylated function had not yet been investigated. We show that multiple prenylated proteins control cardiomyocyte proliferation including RRAS2 and NAP1L4. We demonstrate that prenylation has differential effects on distinct proteins, with RRAS2 prenylation controlling membrane localization and NAP1L4 prenylation regulating cardiomyocyte mitosis and centrosome homeostasis. Together, these data show that protein prenylation is required for cardiomyocyte proliferation through multiple targets and these processes may need to be re-activated for cardiac regeneration.

## Introduction

In the first week of mammalian postnatal life, the maturation of cardiomyocytes (CMs) drives numerous electrophysiological, structural and metabolic changes [1, 2]. During this period, the capacity for CM proliferation is lost, leading to a loss in regenerative potential in adult mammals [3, 4]. Unlike the adult heart, numerous studies have revealed that the neonatal mammalian heart possesses significant regenerative capacity after pathological insult [3, 5–8]. This regeneration is achieved through endogenous CMs undergoing proliferation thereby restoring cardiac muscle and function [3, 9]. Therefore, strategies targeting CM cell cycle re-entry have become a promising approach to recover contractile function after injury.

We previously performed a drug screen in human cardiac organoids (hCOs) to identify compounds capable of inducing CM proliferation, which identified a role for the mevalonate pathway [10]. The mevalonate pathway is an anabolic, metabolic pathway responsible for the synthesis of a range of products including cholesterol, dolichols, and coenzyme Q10 [11, 12] and is primarily controlled by sterol regulatory element-binding transcription factor 2 (SREBF2) [13, 14]. Due to the variety of products derived from the mevalonate pathway, tight regulation exists over the pathway at the transcriptional [15], post-transcriptional [16–18] and post-translational level [19], illustrating the complex nature of mevalonate pathway regulation. In addition to the products derived from the mevalonate pathway, intermediate metabolites such as farnesyl (FPP) and geranylgeranyl pyrophosphate (GGPP) are generated, which are the lipophilic substrates for prenylation [20]. Prenylation is a ubiquitous post-translational modification involving the covalent attachment of either of these intermediates to proteins containing a carboxyl terminal CaaX motif [21]. This hydrophobic modification alters protein function by dictating protein localization to the plasma membrane or membrane-bound organelles as well as regulating bimolecular interactions and protein processing [21].

With respect to the heart, a role for the mevalonate pathway and prenylation has been observed at multiple stages during cardiac development and appears to be conserved between species. Inhibition of farnesylation in mouse embryonic stem cells resulted in impaired primitive streak formation [22] – which is required for mesoderm induction and CM differentiation [23]. In *Drosophila*, upstream inhibition of the mevalonate pathway or geranylgeranylation resulted in defective cardiac development, impaired cardiac function and embryonic lethality [24]. Recently a requirement for the mevalonate pathway in CM proliferation was demonstrated in both hCOs [10] and mice [25], expression of proteins in the pathway are reduced during cardiac maturation coinciding with postnatal CM cell cycle arrest [25–27]. Therefore, the mevalonate pathway could play a key role in cardiomyocyte proliferation and regeneration, yet the downstream mechanisms remain unresolved.

In cancer, it has been shown that prenylation of Ras, Rab and the Rho-family small GTPases are important [28]. Prenylation is required for membrane trafficking and regulation of processes including cytoskeletal regulation, differentiation and proliferation [29]. With respect to CMs, numerous small GTPases have been implicated in cell cycle including CDC42 [30], KRAS [31] and RHOA [32]. However, how the mevalonate pathway and prenylation activate proliferation in cardiomyocytes is still unclear.

In this study, we use a combination of immature human pluripotent stem cell-derived cardiomyocytes (hPSC-CMs) and relatively more mature hCOs as model systems. Using chemical and genetic methods we reveal that prenylation is the key process by which the mevalonate pathway regulates CM proliferation. We identify 40 prenylated proteins in hPSC-CMs using alkyne-tagged prenyl probes and proteomics providing a broad overview of prenylation in CMs. To confirm a potential role in proliferation, we screened a subset of prenylated proteins using siRNA and uncovered that both RRAS2 and NAP1L4 are necessary for CM proliferation. For RRAS2, we show that prenylation is required for membrane localization, a common feature of prenylated Ras proteins. RNA-sequencing confirmed the requirement of NAP1L4 for CM proliferation whilst affinity chromatography mass spectrometry showed that prenylation dictates NAP1L4 protein interactions with a focus on centrosome homeostasis. Finally, we demonstrate that mevalonate supplementation induces CM mitosis without functional consequences in hCOs. Altogether, our study highlights that prenylation is necessary for CM proliferation whilst uncovering multiple prenylated proteins required for human CM proliferation that function through distinct mechanisms.

## Results

### Prenylation is required for hPSC-CM proliferation

To determine specifically how the mevalonate pathway dictates CM proliferation, we used immature, proliferative 2D hPSC-CMs with an active mevalonate pathway. Different branches of the mevalonate pathway were inhibited using four well-studied small molecule inhibitors (red text in Figure 1A). To shut shown the entire pathway, we used simvastatin (SIM), an FDA-approved lipophilic statin that is a competitive, reversible inhibitor of HMGCR thereby preventing all pathway flux [33]. To specifically suppress cholesterol biosynthesis, TAK-475 (TAK) was chosen as it selectively inhibits squalene synthase [34]. To distinguish between the different arms of prenylation, specific inhibitors FTI-277 (FTi) [35] and GGTI-298 (GGTi) [36] were used to prevent protein farnesylation and geranylgeranylation, respectively. After treatment for 3 days (Figure 1B), cells were fixed and stained for cardiomyocytes (α-actinin), general cell cycle (Ki-67) and nuclei (Figure 1C). Each of the four inhibitors resulted in reduced CM nuclei number (Figure 1D) and CM cell cycle activity (Figure 1E) with the most pronounced effects on both parameters observed after inhibition of geranylgeranylation using GGTi (31% and 47% reductions in CM nuclei number and CM cell cycle activity, respectively).

**Figure 1:**
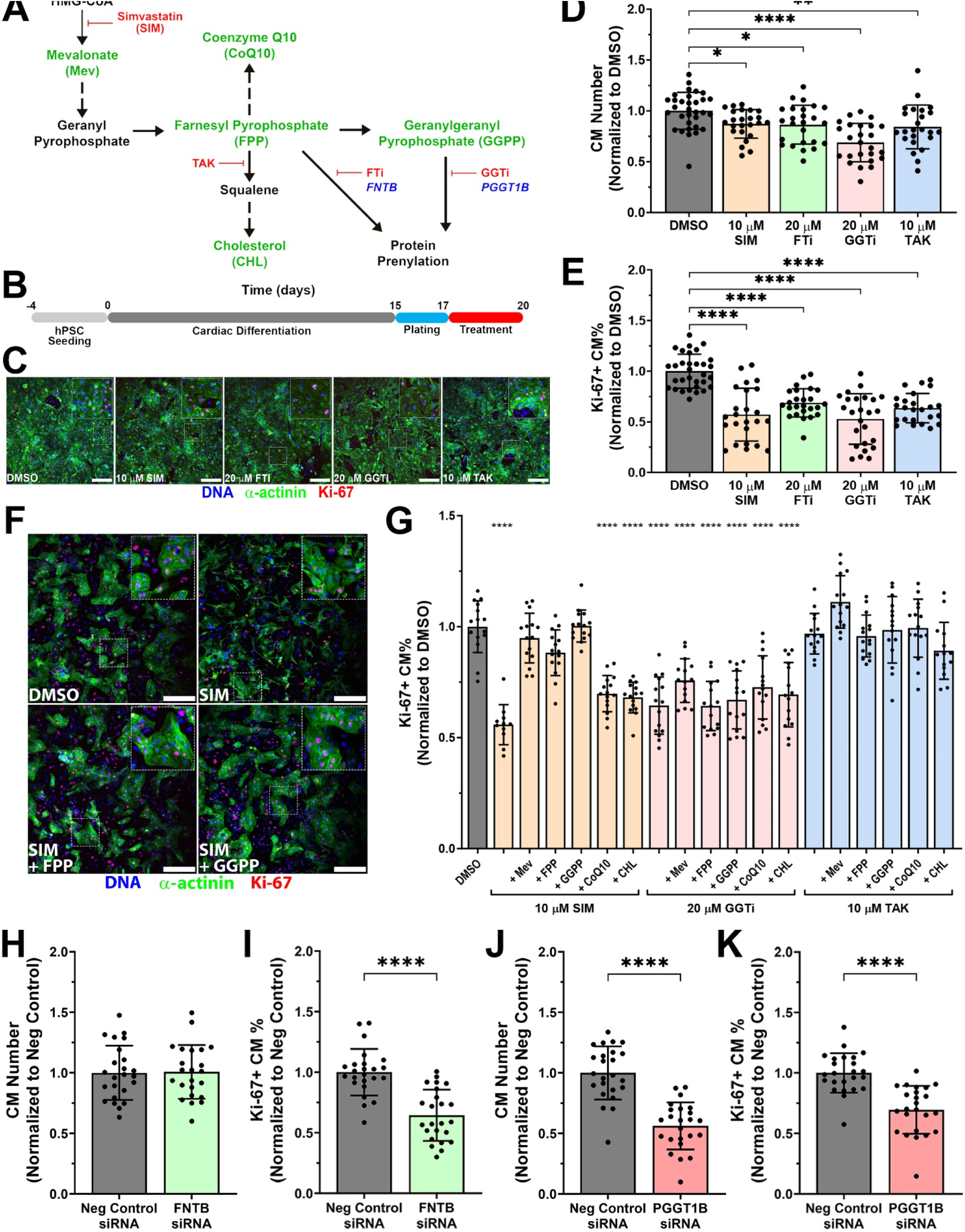
Prenylation components are required for hPSC-CM proliferation. (A) Simple schematic of mevalonate pathway demonstrating intervention points of the four small molecule inhibitors (red text), metabolites used in rescue experiments (green text), and siRNA targets (blue text). (B) Experimental timeline. Immature, proliferative 2D hPSC-CMs were treated for 3 days with small molecule inhibitors. (C) Representative images of 2D hPSC-CMs stained with α-actinin and Ki-67. Scale bar = 250 µm. (D) Each of the four mevalonate pathway inhibitors significantly reduced hPSC-CM nuclei number. n = 24-32 from 3-4 experiments. (E) Each of the four mevalonate pathway inhibitors significantly decreased hPSC-CM cell cycle (Ki-67). n = 24-32 from 3-4 experiments. (F) Representative images of 2D hPSC-CMs from 24-hour rescue experiments stained with α-actinin and Ki-67. Scale bar = 250 µm. (G) Only mevalonate, FPP or GGPP supplementation could rescue hPSC-CM cell cycle in simvastatin-treated cells. GGTi significantly reduced CM cell cycle with no metabolites capable of rescuing cell cycle whilst TAK had no effect on CM cell cycle. n = 11-15 from 3 experiments. (H) FNTB siRNA had no effect on hPSC-CM nuclei number. n = 23-24 from 3 experiments. (I) Knockdown of FNTB significantly reduced hPSC-CM cell cycle. n = 23-24 from 3 experiments. (J) PGGT1B knockdown significantly reduced hPSC-CM nuclei number. n = 24 from 3 experiments. (K) Knockdown of PGGT1B significantly decreased hPSC-CM cell cycle. n = 24 from 3 experiments. Data are presented as mean ± SD. *, **, **** denotes p < 0.05, p < 0.01, p < 0.0001, respectively, compared to DMSO using one-way ANOVA with Dunnett’s post-test (D, E, and G) and compared to Neg Control siRNA using student’s t-test (I-K).

To further delineate the metabolites responsible for CM proliferation, experiments were reduced to 24 hours as metabolites were added exogenously. For these experiments, SIM, GGTi and TAK were used to inhibit different branches of the mevalonate pathway, with pathway-specific metabolites downstream of enzyme inhibition added back to determine if proliferation could be rescued (green text in Figure 1A, Figure 1F). Similar to the 3-day experiments, 24-hour simvastatin treatment resulted in a significant reduction in CM cell cycle activity (Figure 1G). Addition of mevalonate, FPP or GGPP rescued CM cell cycle whereas CoQ10 or cholesterol (CHL) supplementation did not recover CM cell cycle to control levels (Figure 1G). Inhibiting geranylgeranylation also decreased CM cell cycle activity over 24 hours but no metabolites rescued cell cycle under this condition (Figure 1G). Cholesterol biosynthesis inhibition with TAK had no effect on CM cell cycle activity after 24 hours (Figure 1G). This indicated that the reduced CM cell cycle activity observed after 3 days (Figure 1E) may be due to an imbalance in lipid metabolism that takes a longer time to influence proliferation [37]. Collectively, these data demonstrate that prenylation substrates are key mediators of mevalonate-stimulated CM proliferation.

The role of prenylation in controlling CM proliferation was assessed using siRNA. Targeting the beta subunits of either farnesyl or geranylgeranyl prenyltransferase (blue text in Figure 1A), gene knockdown of both *FNTB* and *PGGT1B*, respectively was confirmed using qPCR (Figure S1A). Immature, proliferative 2D hPSC-CMs were transfected with siRNA and cultured for 4 days. *FNTB* knockdown did not change CM nuclei number (Figure 1H), but significantly reduced CM cell cycle activity (Figure 1I). *PGGT1B* resulted in a significant decrease in both CM nuclei number (Figure 1J) and CM cell cycle activity (Figure 1K). These results imply that farnesylation and geranylgeranylation are involved in different stages of cell cycle.

### Proteomics and prenyl probes identify 40 prenylated proteins in hPSC-CMs

To identify prenylated proteins, we first re-analysed our previous hCO proteomic data from our drug screening paper [10]. These analyses revealed two high-confidence farnesylation sites, NAP1L1-C388 (Figure S2A) and NAP1L4-C383 (Figure S2B), that were increased with pro-proliferative compounds (Figure S2C,D). To discover other prenylated proteins that may be involved in hPSC-CM proliferation, we used alkyne-tagged prenyl probes including YnF for farnesylation and YnGG for geranylgeranylation enabling chemical proteomics [38]. These probes were incorporated by hPSC-CMs over 16 hours and the protein lysate harvested and subjected to click chemistry to ligate AzTB – an azide-tagged reagent consisting of a TAMRA fluorophore and a biotin group – allowing in-gel fluorescence detection (Figure 2A). There was a concentration-dependent increase in probe intensity for both YnF and YnGG using in-gel fluorescence analysis (Figure 2B). To demonstrate probe specificity, hPSC-CMs were pre-treated with simvastatin (mevalonate pathway), tipifarnib (farnesylation) or GGTI-298 (geranylgeranylation) for 1 hour before probe addition and subsequent culture for 16 hours.

**Figure 2:**
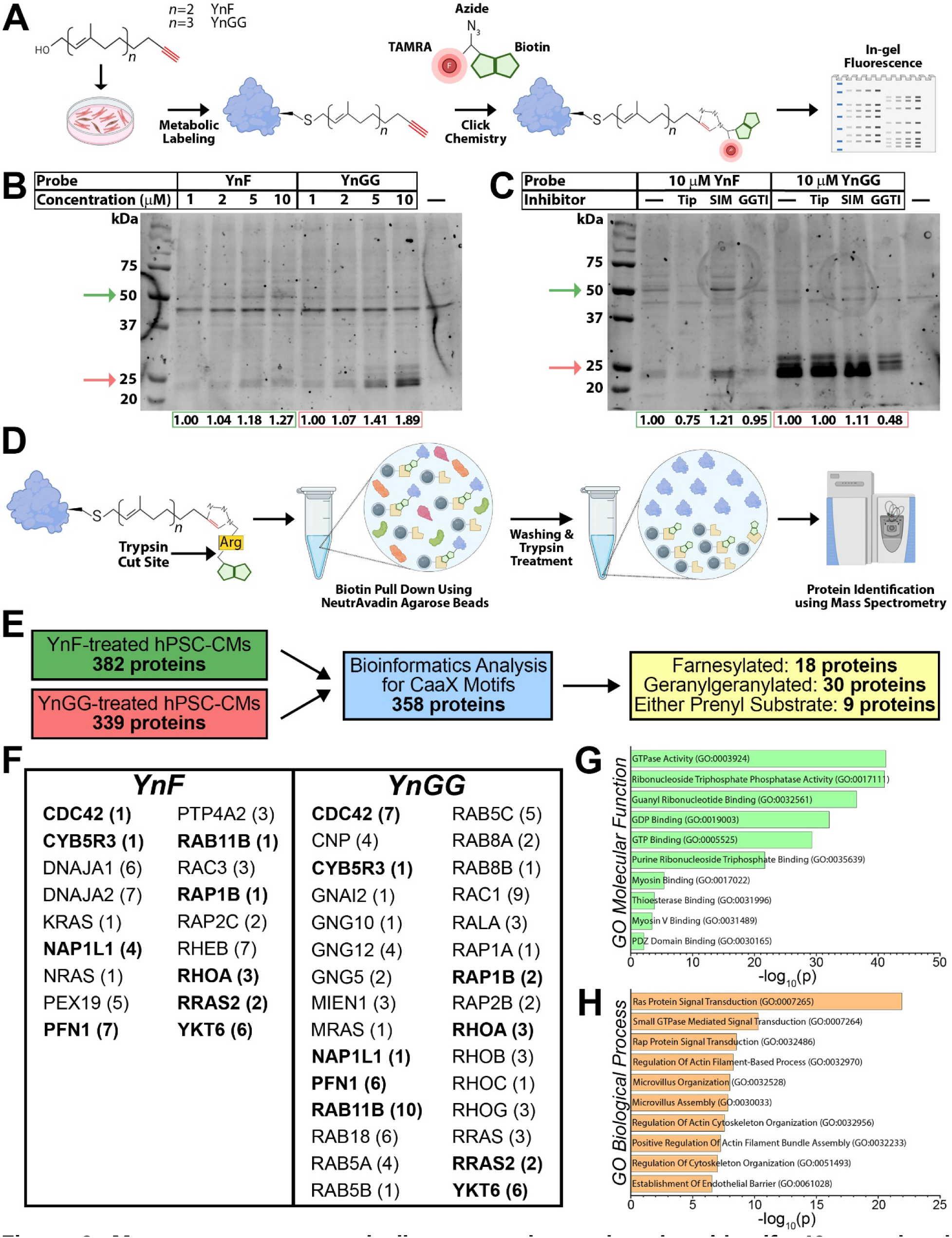
Mass spectrometry and alkyne-tagged prenyl probes identify 40 prenylated proteins in hPSC-CMs. (A) Schematic showing in-gel fluorescence prenyl probe experiments. YnF or YnGG (‘n’ denotes the number of repeats of the structure within the parentheses for each probe) are added directly to cell culture media and cultured for 16 hours. Protein is extracted and subjected to click chemistry where AZTB, an azide-tagged capture reagent with a biotin group and TAMRA fluorophore, binds to the alkyne group in prenyl probe. This can be run on a gel and imaged using a fluorescent imager to assess probe incorporation. (B) Processed lysates from increasing concentrations of YnF and YnGG (1-10 µM) treated hPSC-CMs were run on an SDS-PAGE gel and imaged at 555 nm. Arrows indicate bands used for fluorescence intensity analysis for YnF (green) and YnGG (red) which showed a concentration-dependent increase in band intensity for both YnF (50 kDa) and YnGG (25 kDa). Values are normalized to 1 µM probe. (C) YnF or YnGG (10 µM) were co-cultured with 100 nM tipifarnib (Tipi), 10 µM simvastatin (SIM) or 20 µM GGTi-298 (GGTi) with processed lysate run on SDS-PAGE gel and imaged at 555 nm. Arrows indicate bands used for fluorescence intensity analysis for YnF (green) and YnGG (red). Decreased fluorescence intensity was observed with tipifarnib treatment with YnF at 50 kDa whilst GGTi treatment reduced fluorescence intensity with YnGG at 25 kDa. Simvastatin resulted in increased band intensity using either probe. Values are normalized to 10 µM probe only. (D) Schematic showing prenyl probe experiments for mass spectrometry. The capture reagent for the click chemistry in these experiments was AzRB, an azide-tagged capture reagent containing a biotin group and trypsin cut site. Processed lysate was incubated with NeutrAvidin beads to allow biotin-streptavidin binding followed by a series of washing steps to remove unbound proteins. Trypsin treatment allowed release of probe-labelled peptides into supernatant and subsequent identification by mass spectrometry. (E) Analysis pipeline illustrating the cross-referencing of mass spectrometry-identified proteins in YnF and YnGG-treated hPSC-CMs with CaaX motif-containing proteins generated through bioinformatics enabling the discovery of prenylated proteins expressed by hPSC-CMs. (F) Mass spectrometry identified 18 farnesylated and 30 geranylgeranylated proteins with 9 proteins (bold text) modified by either prenyl substrate. The number of unique peptides detected for each protein are indicated in parentheses. n = 1 experiment. (G) Enrichr processing of 40 prenylated proteins identified by mass spectrometry and chemical proteomics revealed GTPase activity as most significant term relating to molecular function. (H) Enrichr processing of same list identified Ras protein signaling transduction as most upregulated annotation with respect to biological process.

YnF-treated hPSC-CMs reduced probe fluorescence upon tipifarnib treatment, consistent with impaired farnesyltransferase activity, whilst simvastatin increased fluorescence due to a reduction in endogenous prenyl substrate availability (Figure 2C). YnGG fluorescence was greater than YnF however there was still a subtle increase in fluorescence intensity upon simvastatin treatment. Consistent with GGTI-298 inhibiting geranylgeranylation, there was lower YnGG fluorescence observed (Figure 2C).

To identify prenylated proteins expressed in hPSC-CMs, we performed qualitative chemical proteomics using the same 16-hour incubation. For this, click chemistry ligation was performed using AzRB – an azide-tagged reagent containing a biotin group to assist protein pulldown and a trypsin-cleavable sequence to enhance the release and detection of modified proteins upon digestion (Figure 2D). After click chemistry, lysates were incubated with NeutrAvidin beads to pulldown biotin-tagged proteins before trypsin treatment allowed isolation of modified peptides which were analysed by mass spectrometry (Figure 2D). To detect enriched proteins, probed hPSC-CMs incubated with no AzRB used during click chemistry were used as a negative control. Mass spectrometry and subsequent analysis identified 382 and 339 proteins in YnF and YnGG-treated hPSC-CMs, respectively (Table S1,2). To confirm which of these proteins are capable of being prenylated, bioinformatics analysis was undertaken to identify proteins containing a carboxyl-terminal CaaX motif. To do this, the NCBI RefSeq Protein database (GRCh38) was used to filter for proteins that contained a cysteine at the 4th last residue and prenylation-specific amino acids in the “aX” positions based on a study that performed crystallographic modelling of prenyltransferases to identify CaaX motifs [39]. After protein isoforms were removed, this provided a list of 358 potentially prenylated proteins (Figure S3A, Table S3). Cross-referencing between the mass spectrometry-identified and bioinformatics protein lists (Figure 2E) revealed 39 prenylated proteins in total – 18 farnesylated and 30 geranylgeranylated, with 9 proteins prenylated with either substrate (Figure 2F). Within this list of 39 prenylated proteins, only CDC42 and PFN1 were present in the unprobed control, however compared to YnF and/or YnGG protein coverage was much lower (Figure S3B,C). With respect to PFN1, this is the first analysis to identify it as prenylated protein with enrichment in both YnF and YnGG samples (Figure S3B,C). Combining our data which identified NAP1L4 as a farnesylated protein in hCOs with this new dataset, this provided a list of 40 prenylated proteins in hPSC-CMs. When this list was processed through Enrichr [40–42], the gene ontologies for molecular function (Figure 2G) and biological process (Figure 2H) showed an enrichment for GTPase activity and Ras protein signalling, respectively (Table S4,5). Collectively, these findings characterize prenylated proteins within hPSC-CMs and provide a target list of prenylated proteins to allow interrogation of their potential role in CM proliferation.

### NAP1L4 and RRAS2 are required for hPSC-CM proliferation

After observing the increase in abundance of farnesylated NAP1L1 and NAP1L4 after pro-proliferative treatment, we first wanted to determine if NAP1L1 or NAP1L4 were directly involved in hPSC-CM proliferation. Both proteins are responsible for assembling nucleosomes by binding histones, albeit with differing affinities [43, 44], and transferring them to DNA [43]. NAP1L1 and NAP1L4 have been shown to be prenylated elsewhere [38], but their role in hPSC-CM proliferation and impact of prenylation are currently undefined.

2D hPSC-CMs were transfected with siRNA targeting NAP1L1 or NAP1L4 and cultured for 4 days, with gene knockdown confirmed using qPCR (Figure S1B). Silencing of NAP1L1 had no effect on hPSC-CM nuclei number (Figure 3A) however significantly reduced hPSC-CM cell cycle activity (Figure 3B). Conversely, knockdown of NAP1L4 decreased both hPSC-CM nuclei number (Figure 3C) and hPSC-CM cell cycle activity (Figure 3D). This indicates that NAP1L4 plays a key role in hPSC-CM proliferation.

**Figure 3:**
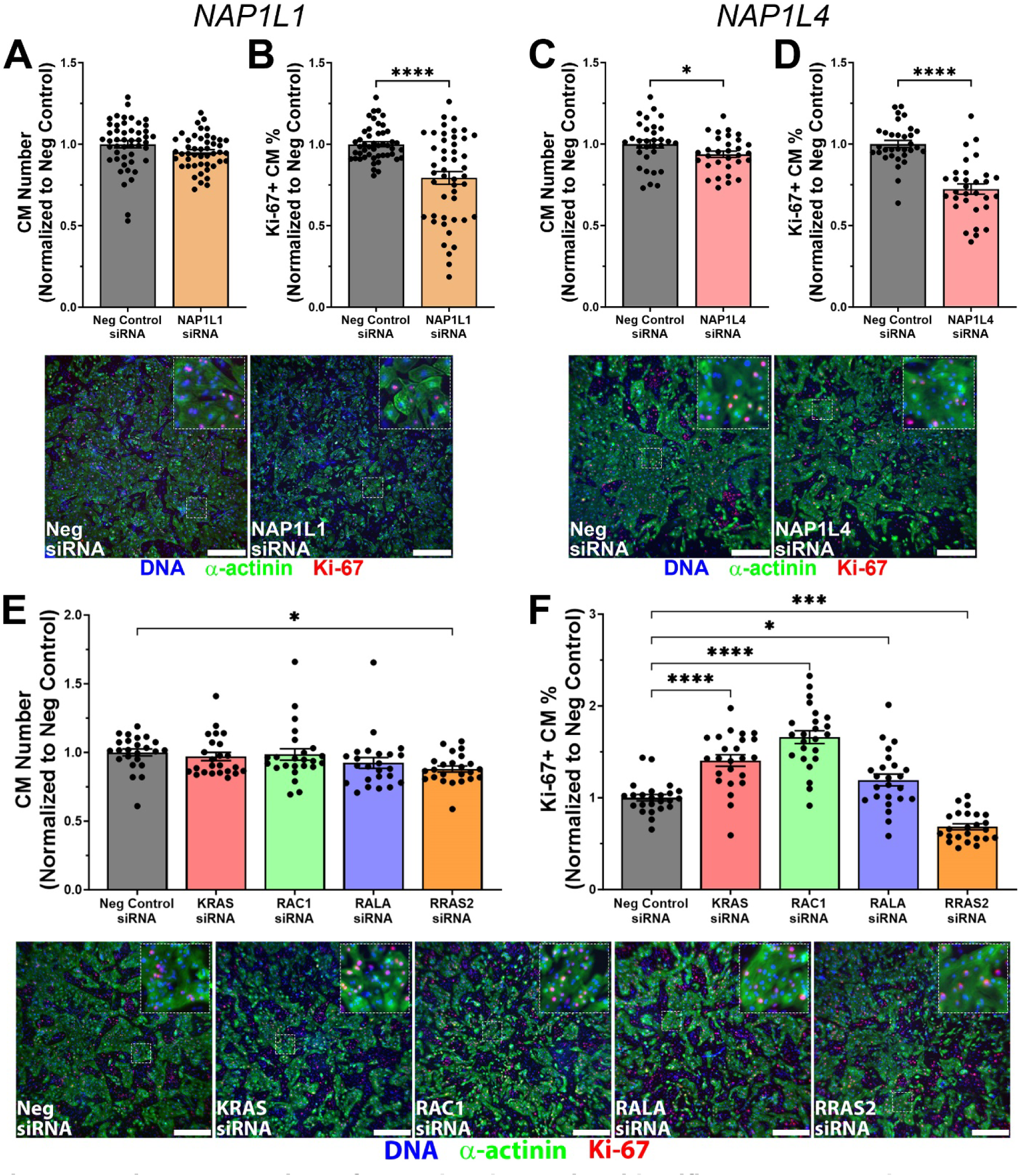
siRNA screening of prenylated proteins identifies NAP1L4 and RRAS2 as requirements for hPSC-CM proliferation. (A) NAP1L1 knockdown had no effect on hPSC-CM nuclei number. n = 47-48 from experiments. (B) Knockdown of NAP1L1 significantly reduced hPSC-CM cell cycle activity. Representative images of α-actinin and Ki-67-stained NAP1L1 siRNA transfected immature hPSC-CMs. Scale bar = 500 µm. n = 47-48 from 6 experiments. (C) Silencing of NAP1L4 significantly decreased hPSC-CM nuclei number. n = 31-32 from 4 experiments. (D) NAP1L4 knockdown significantly reduced hPSC-CM cell cycle activity. Representative images of α-actinin and Ki-67-stained NAP1L4 siRNA transfected immature hPSC-CMs. Scale bar = 500 µm. n = 31-32 from 4 experiments. (E) Silencing of RRAS2 significantly reduced CM number. n = 24 from 3 experiments. (F) Knockdown of KRAS, RAC1 and RALA significantly elevated the number of Ki-67+ CMs whilst RRAS2 silencing decreased CM cell cycle. Representative images of siRNA transfected 2D hPSC-CMs stained with α-actinin (green) and Ki-67 (red). Scale bar = 500 µm. n = 24 from 3 experiments. Data are presented as mean ± SEM. *, ***, **** denotes p < 0.05, p < 0.001, p < 0.0001, respectively, compared to Neg Control siRNA using student’s t-test (A-D) or using one-way ANOVA with Dunnett’s post-test (E-F).

Given that Ras proteins are involved in cell proliferation [45], a subset of Ras signaling proteins (RRAS2, KRAS, RAC1 and RALA) were chosen for a siRNA screen to assess their role in CM cell cycle (Table S5). Firstly, we confirmed knockdown of target gene expression using qPCR (Figure S1C). With respect to hPSC-CM cell cycle, when KRAS, RAC1 and RALA were silenced there was no change in hPSC-CM nuclei number (Figure 3E) with significant increases in cell cycle (Figure 3F). This potentially indicates a hypertrophic response. Conversely, RRAS2 silencing resulted in a significant drop in both hPSC-CM nuclei number (Figure 3E) and cell cycle (Figure 3F).

Overall, the hPSC-CM cell cycle activity decreases of 28% and 32% in NAP1L4 and RRAS2 knockdown, respectively, are similar to the 30% with PGGT1B knockdown. Given this is less effective than small molecule inhibition, this potentially indicating that multiple prenylation processes control proliferation or residual activity remains after knockdown. Together, this indicates that two prenylated proteins, NAP1L4 and RRAS2, are critical hPSC-CM cell cycle regulators.

### RNA-sequencing indicates that NAP1L4 knockdown affects the transcription of cell cycle genes

To gain insights into how NAP1L4 is involved in hPSC-CM proliferation, we performed bulk RNA-sequencing on 2D hPSC-cardiac cells from two cell lines (HES3 and PGP1) subjected to NAP1L4 siRNA for two days. NAP1L4 knockdown was confirmed (Figure S4A). Principal component analysis (PCA) separated the cell lines on PC1, likely due to non-myocyte differences (Figure 4A). The negative control and NAP1L4 siRNA-treated hPSC-CMs were clearly separated on PC2 for both cell lines (Figure 4A). There were 1478 differentially expressed genes (DEGs) in HES3 and 2071 DEGs in PGP1 (DEGs, FDR < 0.05 with a minimum log2-fold-change of |1.2|). Of these, 520 were commonly up-regulated and 523 were commonly down-regulated across both cell lines in NAP1L4 versus negative control siRNA (Figure S4B, S4C, Table S6 – 8). Gene Ontology analysis of the common down-regulated genes resulted in enrichment of terms related to cardiac muscle contraction, cardiac muscle development and DNA replication initiation (Figure 4B, Table S9). Specifically looking at cell cycle genes, there was reduced expression of genes from G1 (*CCNE2, CDC6, CDK16, ESRRB, SASS6*), S (*ATRX*), and G2/M (*ANLN, CCNA2, CDC14B, CDC25A, CENPE*) phases (Figure 4C). Together, these results reveal that knockdown of NAP1L4 can impact the transcription of cell cycle genes, potentially via its nuclear chaperone role.

**Figure 4:**
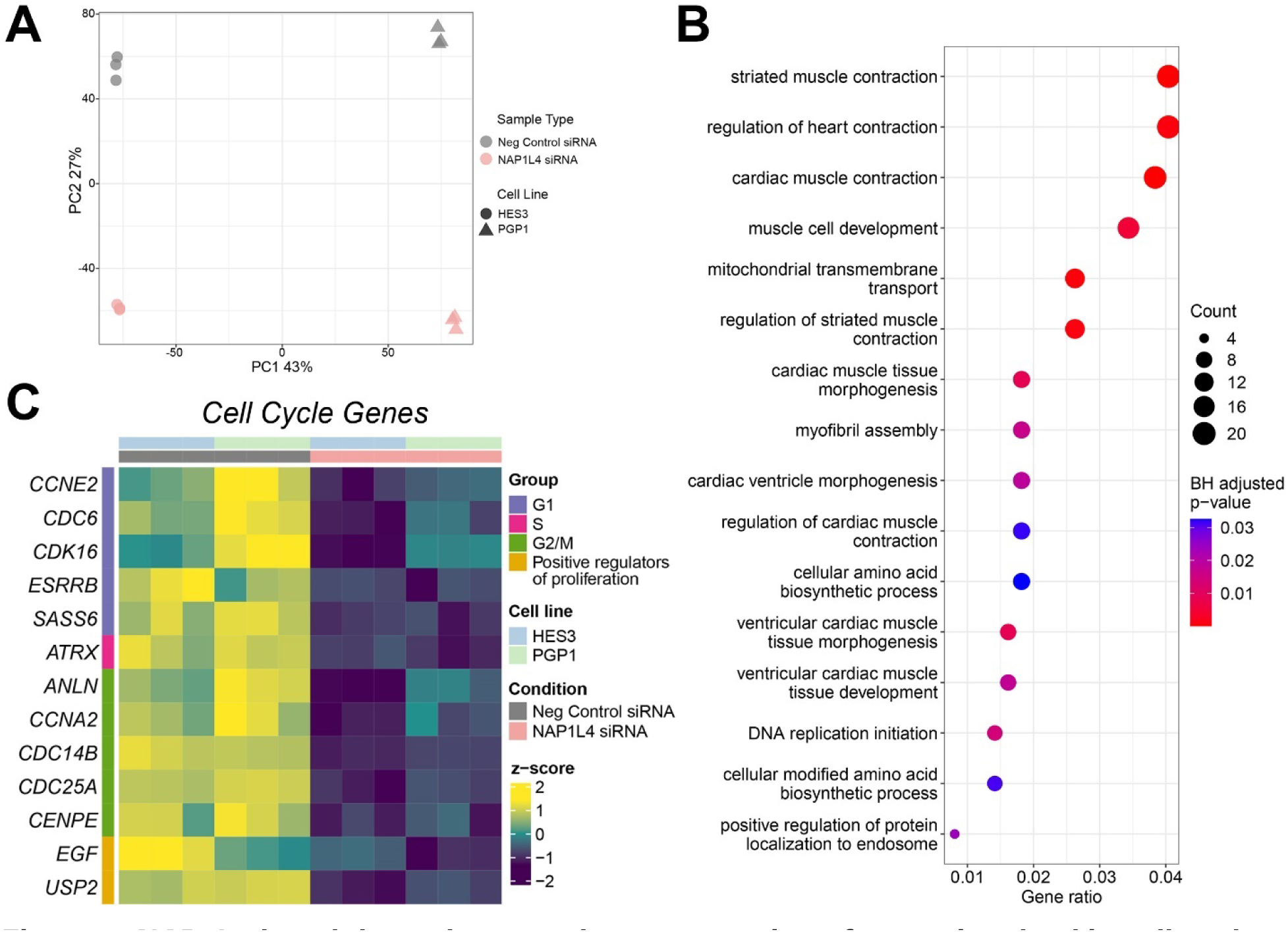
NAP1L4 knockdown downregulates expression of genes involved in cell cycle. (A) PCA of protein coding genes for 2D hPSC-CMs subjected to NAP1L4 siRNA or Neg Control siRNA for HES3 and PGP1 cell lines. n = 3 from 1 experiment per cell line. (B) Top gene ontology terms from sub-ontology biological process enriched based on common down-regulated differentially expressed genes across HES3 and PGP1 cell lines in NAP1L4 siRNA-treated 2D hPSC-CMs compared to Neg Control siRNA. (C) Heatmap illustrating significantly down-regulated cell cycle genes in NAP1L4 siRNA-treated 2D hPSC-CMs compared to Neg Control siRNA across HES3 and PGP1 cell lines.

### NAP1L4 overexpression alone is insufficient to activate cell cycle

To determine the pro-proliferative capacity of NAP1L4 using overexpression, we used our hCO maturation protocol. This drives advanced maturation at the level of transcription, function, metabolism and cell cycle [46]. NAP1L4 overexpression (or a GFP control) was targeted to cardiomyocytes using delivery of adeno-associated virus 6 (AAV6) with a CM-specific cardiac troponin T promoter for 4 days (Figure S5A). Overexpression of NAP1L4 did not impact contractile function (Figure S5B). NAP1L4 overexpression also did not alter whole hCO Ki-67 intensity (Figure S5C), or mitosis marked by phospho-histone H3-positive (pHH3+) CMs undergoing sarcomere disassembly (Figure S5D). This indicates that NAP1L4 itself is not sufficient to drive hPSC-CM proliferation. This is perhaps due to insufficient prenylation or that other prenyl-controlled proteins such as RRAS2 may by synergistic activators of cell cycle.

### Mevalonate supplementation induces cardiomyocyte mitosis in human cardiac organoids

Expression of key genes in the mevalonate pathway are controlled by SREBF2 which are repressed during cardiac maturation [26]. To determine if mevalonate pathway gene overexpression is sufficient to induce hPSC-CM proliferation, 2D hPSC-CMs were cultured in maturation medium which reduces but does not arrest cell cycle in 2D culture. They were then transfected with a modified mRNA (modRNA) to overexpress an active form of SREBF2 (mSREBF2). mSREBF2 is the transcriptionally active, processed form of SREBF2 which occurs upon cleavage at the Golgi by resident site-1 and site-2 proteases [47] (Figure S6A). We firstly confirmed sustained overexpression in 75% of hPSC-CM after 3 days using 1 µg eGFP modRNA (Figure S6B). qPCR confirmed increases in multiple SREBF2 target genes after 4 hours (Figures S6A,C). However, 2 days of mSREBF2 modRNA treatment led to reduced CM proliferation (Figure S6D).

After observing that transcriptional activation of mevalonate pathway genes was insufficient to drive hPSC-CM proliferation, we wanted to determine if we could instead increase metabolic substrates. To do this, mature hCOs were directly supplemented with 500 µM mevalonate for 2 days (Figure 5A). Mevalonate addition had no effect on contractile force (Figure 5B) or rate (Figure 5C). However, there were significant reductions in the 50% activation time (Figure 5D) and 50% relaxation time (Figure 5E). These >5% alterations in the absence of a rate change indicate improved contractile kinetics which has been associated with improved metabolic function [48–51]. Mevalonate treatment significantly increased whole-tissue Ki-67 intensity (Figure 5F) accompanied by an increase in mitosis marked by phospho-histone H3-positive (pHH3+) CMs undergoing sarcomere disassembly (Figure 5G). Collectively, these results show that direct supplementation of the upstream metabolite of the mevalonate pathway is sufficient to improve contractile kinetics and induce hPSC-CM mitosis. This indicates that metabolic substrate provision is a more effective target than pathway enzyme abundance to induce proliferation, and induction of prenylation and its downstream effects on proteins might be a useful avenue for activation of proliferation. We therefore next determined the role of prenylation for the critical proliferation regulators RRAS2 and NAP1L4.

**Figure 5:**
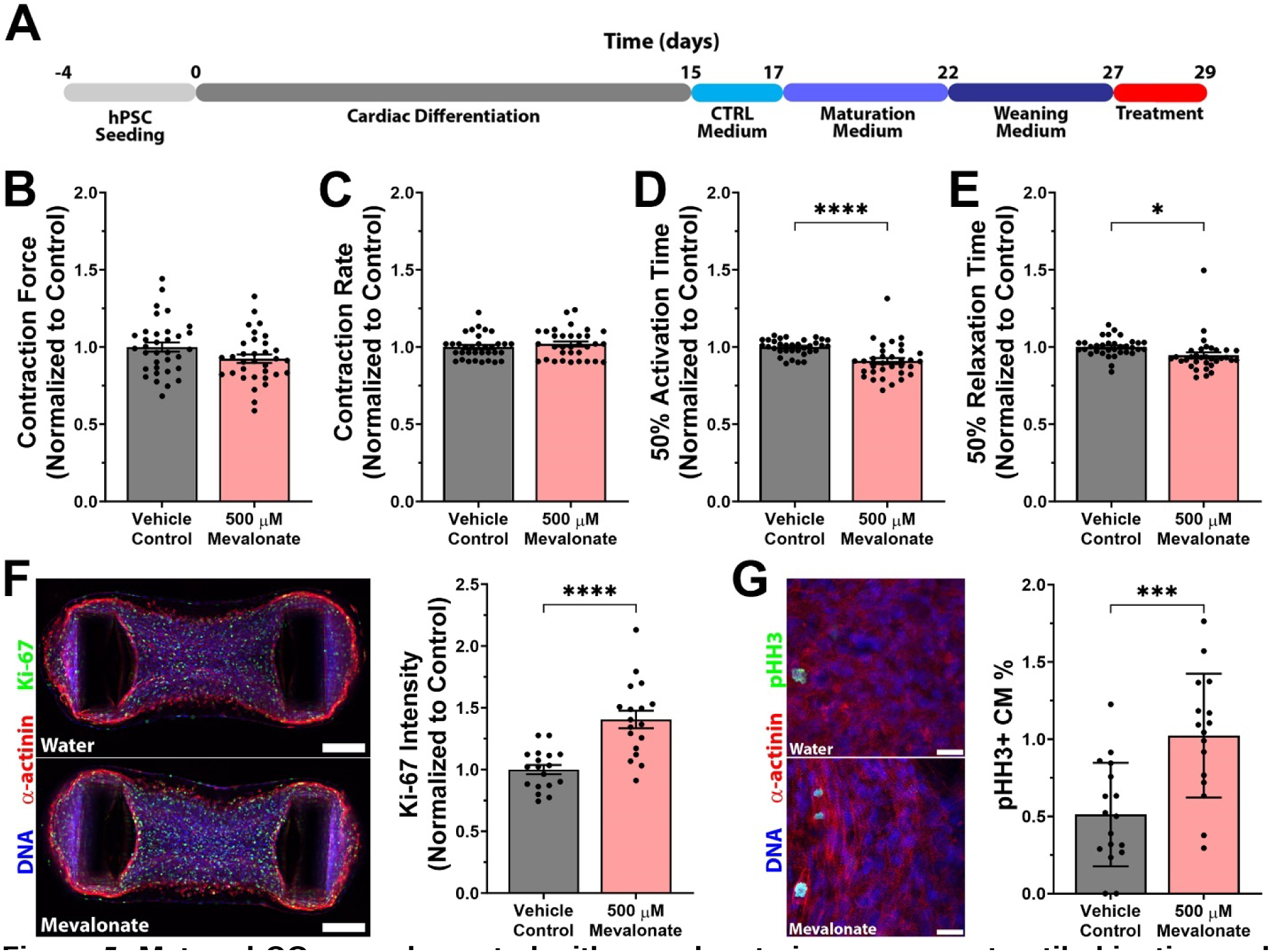
Mature hCOs supplemented with mevalonate improves contractile kinetics and significantly increases CM mitosis. (A) Schematic of experimental pipeline. Mature hCOs were supplemented with mevalonate (500 µM) for 2 days. (B) Mevalonate treatment had no effect on contractile force. n = 33-34 hCOs from 3 experiments. (C) Mevalonate supplementation did not alter contractile rate. n = 33-34 hCOs from 3 experiments. (D) Mevalonate treatment significantly decreased 50% activation time. n = 33-34 hCOs from 3 experiments. (E) Mevalonate supplementation significantly reduced 50% relaxation time. n = 33-34 hCOs from 3 experiments. (F) Representative images of immunostained hCOs used for whole tissue intensity analysis for Ki-67. Scale bar = 200 µm. Mevalonate supplementation significantly increased Ki-67 intensity. n = 18 hCOs from 3 experiments. (G) Representative images of pHH3-stained hCOs to quantify CM mitosis. Scale bar = 20 µm. Mevalonate treatment significantly increased CM mitosis (pHH3+ CMs). n = 16-17 hCOs from 3 experiments. Data are presented as mean ± SEM (B-F) or ± SD (G). *, ***, **** denotes p < 0.05, p < 0.001, p < 0.0001, respectively, compared to vehicle control using student’s t-test.

### RRAS2 prenylation controls membrane localization

As Ras proteins elicit diverse signaling responses from different complexes and locations within the cell [52], we wanted to determine the reliance of RRAS2 localization on prenylation in hPSC-CMs. To do this, we synthesized a RRAS2 modRNA with an N-terminal human influenza hemagglutinin (HA) tag, confirmed by agarose gel electrophoresis (Figure S7A,B). hPSC-CMs were treated with simvastatin or DMSO vehicle for 1 hour before being transfected with modRNA (Figure 6A). After 24 hours, RRAS2 was localized primarily at the plasma membrane and was localized to the nucleus in the simvastatin treated group (Figure 6B). This is consistent with the classical requirement of prenylation for Ras protein docking into the membrane for activation.

**Figure 6:**
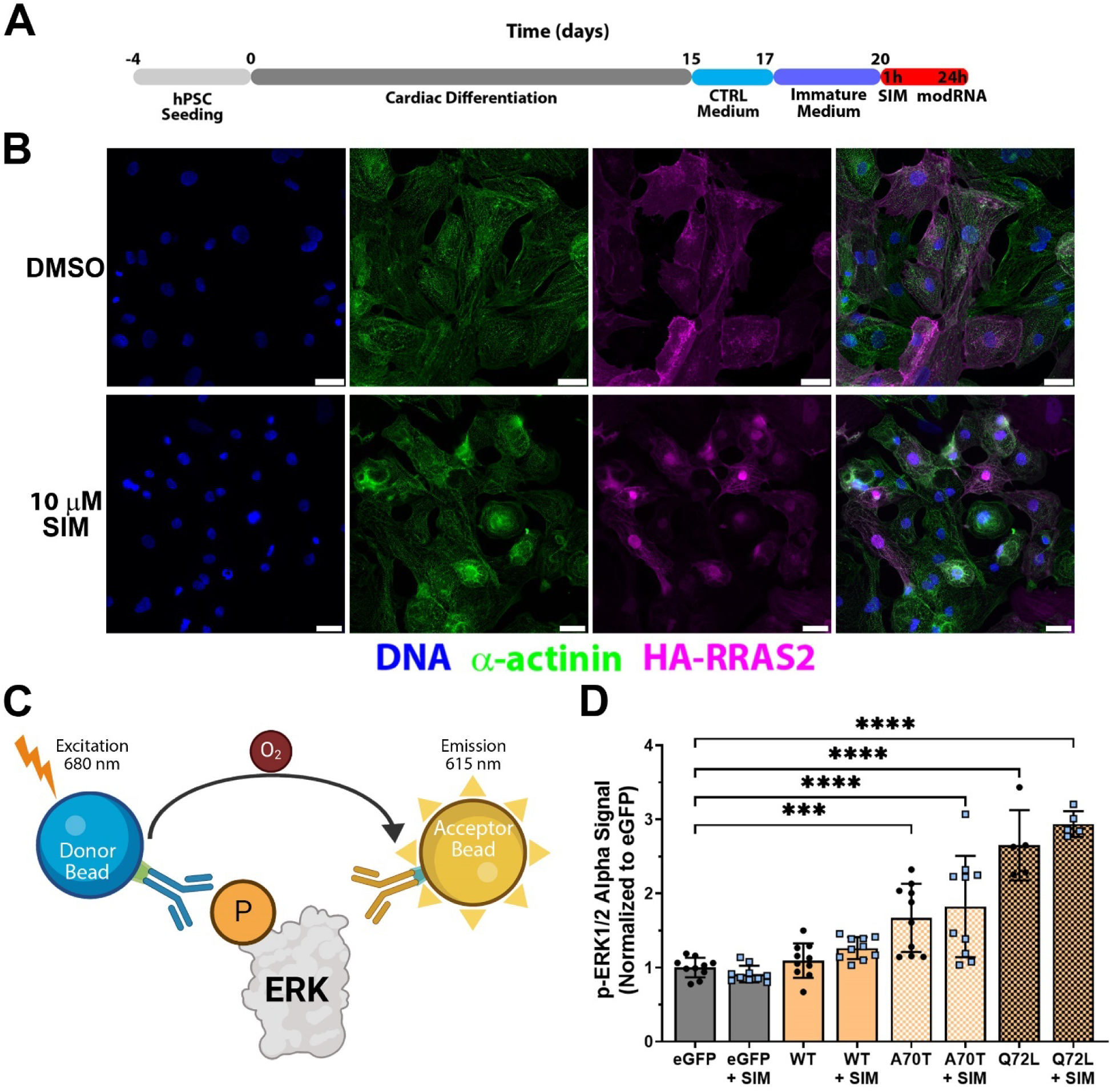
RRAS2 prenylation is required for plasma membrane localisation in hPSC-CMs. (A) Experimental timeline. Immature, proliferative 2D hPSC-CMs were cultured for 1 hour with simvastatin before being transfected with HA-tagged modRNA for 24 hours. (B) Immunofluorescence images showing that RRAS2 (magenta) is localized at the plasma membrane in hPSC-CMs (green) however simvastatin treatment results in localization in the nucleus (blue). Scale bar = 25 µm. (C) Schematic of the AlphaLISA Surefire Ultra assay. When excited with light, the donor bead releases a free oxygen molecule which allows the acceptor bead to emit light which is directly proportional to the abundance of phosphorylated protein. (D) Overexpression of either RRAS2 mutant (A70T or Q72L) significantly elevated p-ERK1/2 which was unaffected by co-treatment with simvastatin. n = 5-10 from 1-2 experiments. Data are presented as or mean ± SD. ***, **** denotes p < 0.001, p < 0.0001, respectively, compared to eGFP using one-way ANOVA with Dunnett’s post-test.

As extracellular signal-regulated kinase (ERK) is classically downstream from Ras proteins [53], we performed a phosphorylated ERK assay to assess potential impact on cellular signaling. To do this, we used the AlphaLISA phospho-ERK1/2 (p-ERK1/2) assay (Figure 6C). In these assays, we inhibited prenylation using simvastatin for 1 hour before modRNA transfection and used activating RRAS2 mutants A70T and Q72L [54] as positive controls. We found that ERK phosphorylation was not dependent on prenylation, with only mutant RRAS2 capable of inducing large increases in pERK (Figure 6D). This indicates that RRAS2 activation of ERK may require additional signaling drivers for proliferation or that it may be working via alternative signaling networks which will be interesting to follow-up in the future.

### Prenylation of NAP1L4 controls mitotic transition

To better understand the function of NAP1L4 in a dynamic setting, live-cell imaging was performed. To do this, we synthesized a modRNA encoding NAP1L4 with eGFP fused to the N-terminus. To determine if prenylation alters NAP1L4 localization, we synthesized a prenyl mutant where C383 in the CaaX motif was substituted with a serine to prevent farnesylation (NAP1L4-C383S) (Figure S7C). After confirming that GFP-NAP1L4 modRNA was the correct molecular size (Figure S7D), we transfected 2D hPSC-CMs for 8 hours before performing live-cell imaging for 20 hours. GFP-NAP1L4 entered the nucleus at the start of mitosis and stayed in the nucleus until karyokinesis completion where it then translocated back to the cytoplasm (Figure 7A). Conversely, GFP-NAP1L4-C383S also entered the nucleus upon initiation of mitosis, but then the cell stalls and ultimately leaves the nucleus after a prolonged period with karyokinesis only completed 33% of the time (Figure 7A,B). Aside from incomplete karyokinesis, quantification of time spent in the nucleus revealed GFP-NAP1L4-C383S spent twice as much time in the nucleus compared to GFP-NAP1L4, 286 versus 143 minutes, respectively (Figure 7C).

**Figure 7:**
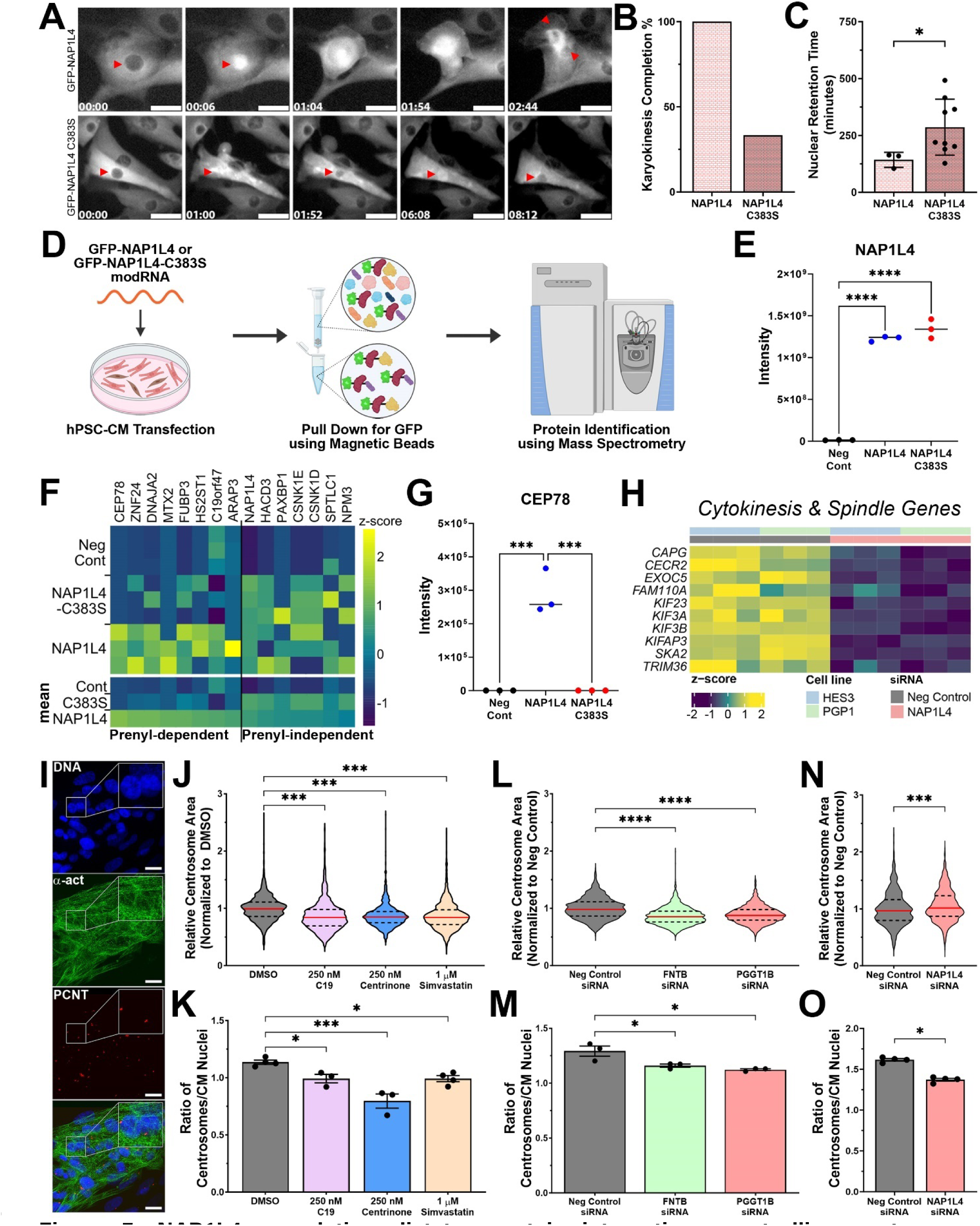
NAP1L4 prenylation dictates protein interactions controlling centrosome duplication and mitosis. (A) Still images of GFP-NAP1L4 and GFP-NAP1L4-C383S during cell division. GFP-NAP1L4 enters the nucleus (red arrowhead) at the initiation of division and exits upon karyokinesis completion. GFP-NAP1L4-C383S enters the nucleus (red arrowhead) however mitosis is delayed and/or aborted and ultimately NAP1L4 exits the nucleus. Time scale = hours:minutes. Scale bar = 20 µm. (B) Upon initiation of mitosis, entry of GFP-NAP1L4-C383S into the cell resulted in complete cytokinesis only 33% of the time whereas GFP-NAP1L4 completed cytokinesis 100% of the time. n = 3-9 nuclei from 2 experiments. (C) Total nuclear retention time doubled for GFP-NAP1L4-C383S compared to GFP-NAP1L4 during cell division. n = 3-9 nuclei from 2 experiments. (D) Schematic of experimental pipeline. 2D hPSC-CMs cultured in maturation media were transfected with GFP-NAP1L4 or GFP-NAP1L4-C383S modRNA for 8 hours before lysis. Protein lysate was subjected to pulldown using GFP-targeting magnetic beads which was subsequently cleaved, processed and analyzed using mass spectrometry. (E) Raw intensity values from protein pulldown for NAP1L4. n = 3 from 1 experiment. (F) Heatmap highlighting proteins enriched by prenylation (delta Z-score > 1 between GFP-NAP1L4 and GFP-NAP1L4-C383S). n = 3 from 1 experiment. (G) Raw intensity values from protein pulldown for CEP78. n = 3 from 1 experiment. (H) Heatmap illustrating significantly down-regulated genes from RNA-sequencing experiment associated with cytokinesis or mitotic spindle in NAP1L4 siRNA-treated 2D hPSC-CMs compared to Neg Control siRNA across HES3 and PGP1 cell lines. (I) Representative image showing 2D hPSC-CMs stained with α-actinin (α-act) and centrosomal marker, pericentrin (PCNT). Scale bar = 20 µm. (J) Simvastatin treatment significantly reduced centrosomal area in CMs. n = 1002-1715 centrosomes from 3-4 experiments. (K) Simvastatin decreased number of centrosomes per CM nuclei. n = 3-4 experiments (which is from 1002-1715 centrosomes total and 1012-1672 CM nuclei). (L) Silencing of FNTB or PGGT1B resulted in decreased centrosomal area in CMs. n = 1245-1501 centrosomes from 3 experiments. (M) FNTB and PGGT1B knockdown also reduced number of centrosomes per CM nuclei. n = 3 experiments (which is from 1245-1501 centrosomes total and 967-1311 CM nuclei total). (N) NAP1L4 knockdown significantly increased hPSC-CM centrosomal area. n = 750-780 centrosomes from 4 biological replicates across 2 experiments. (O) NAP1L4 siRNA significantly decreased the number of centrosomes per hPSC-CM nuclei. N = 4 biological replicates across 2 experiments (from 750-780 centrosomes total and 464-568 hPSC-CM nuclei in total). Data are presented as mean ± SD (C), median with interquartile range (J, L, N) or mean ± SEM (K, M, O). *, ***, **** denotes p < 0.05, p < 0.001, p < 0.0001, respectively compared to NAP1L4 WT (C), DMSO (J, K) or Neg Control siRNA (L-O) using unpaired t-test (C), one-way ANOVA with Dunnett’s post-test (J-M), student’s t-test (N) or unpaired t-test with Mann Whitney test (O).

### NAP1L4 prenylation dictates centrosome interactions

After observing that prenylation impacted the mitotic transition, we next sought to determine which proteins interacted with NAP1L4 and whether prenylation alters these interactions. To do this we pulled down NAP1L4 and NAP1L4-C383S using the N-terminal eGFP tag. 2D hPSC-CMs were transfected with GFP-NAP1L4 or GFP-NAP1L4-C383S for 8 hours, prior to lysis, pulldown and mass spectrometry (Figure 7D). We firstly confirmed NAP1L4 only in the modRNA groups (Figure 7E). We also found that negative controls (RNAiMAX only) were distinctly separated from GFP-NAP1L4 and GFP-NAP1L4-C383S in a PCA, with a subtle separation between GFP-NAP1L4 and GFP-NAP1L4-C383S (Figure S8). There were a core set of proteins shared by both GFP-NAP1L4 and GFP-NAP1L4-C383S (Figure 7F), indicating a prenyl-independent role consistent with its reported canonical function as a nuclear chaperone. These interactions include CSNK1D and CSNK1E, which are the casein kinases responsible for NAP1L4 trafficking to the nucleus [55]. Also PAXBP1, an indispensable DNA-binding protein required for S-phase entry [56], and NPM3, which also functions as a histone chaperone [57].

With respect to prenyl-dependent interactions, there was an enrichment for 8 proteins in GFP-NAP1L4 over GFP-NAP1L4-C383S, including the interaction with CEP78 that was completely abolished in the GFP-NAP1L4-C383S prenyl mutant (Figure 7G). CEP78 is responsible for regulating centrosome homeostasis [58], so we investigated the effect of NAP1L4 knockdown on cytokinesis and mitotic spindle gene expression. Genes from both processes had reduced expression in NAP1L4 siRNA-treated hPSC-CMs (Figure 7H, Table S10), but did not include the interacting proteins themselves.

This new relationship led us to explore whether the mevalonate pathway influences centrosome duplication in hPSC-CMs. To do this, 2D hPSC-CMs were treated with simvastatin for 3 days and compared with two positive controls: centrinone (PLK4 inhibitor) and C19 (MST2 activator) [59]. After treatment, hPSC-CMs were stained for the centrosomal marker, pericentrin, and centrosome area and number were quantified in hPSC-CMs (Figure 7I). Inhibition of the mevalonate pathway reduced both centrosome area (Figure 7J) and number per CM nuclei (Figure 7K) similar to the positive controls. To determine if this was a prenylation-specific mechanism, 2D hPSC-CMs were transfected with siRNA targeting FNTB and PGGT1B and cultured for 4 days. Silencing of farnesylation or geranylgeranylation significantly decreased centrosome area (Figure 7L) and number per CM nuclei (Figure 7M). To validate if a functional relationship existed between NAP1L4 and hPSC-CM centrosome homeostasis, we treated 2D hPSC-CMs with NAP1L4 siRNA for 4 days. NAP1L4 knockdown resulted in an increase in CM centrosome area (Figure 7N) and a significant decrease in centrosome number per CM nuclei (Figure 7O). This relationship between NAP1L4 and centrosome duplication is not a general prenylated protein or proliferation effect, as RRAS2 knockdown had no effect on either centrosome area (Figure S9A) or centrosome number per CM nuclei (Figure S9B). These findings indicate that NAP1L4 prenylation plays a specific role in centrosome homeostasis distinct from its nuclear chaperone role.

## Discussion

Targeted induction of CM proliferation to promote endogenous regeneration would be transformative for the treatment of heart failure. In our previous drug screen in mature hCOs, we revealed that the responses for multiple drugs converged on the mevalonate pathway to induce hPSC-CM proliferation [10]. In this study, we provide mechanistic insight into how the mevalonate pathway drives hPSC-CM proliferation (Figure 8). We demonstrate that 1) prenylation is involved in proliferation, 2) prenylated proteins NAP1L4 and RRAS2 are required for cell cycle, 3) NAP1L4 and RRAS2 prenylation dictate protein-protein interactions and protein localization, respectively, and 4) NAP1L4 prenylation regulates interactions with a specific role in centrosome homeostasis and mitotic progression.

**Figure 8:**
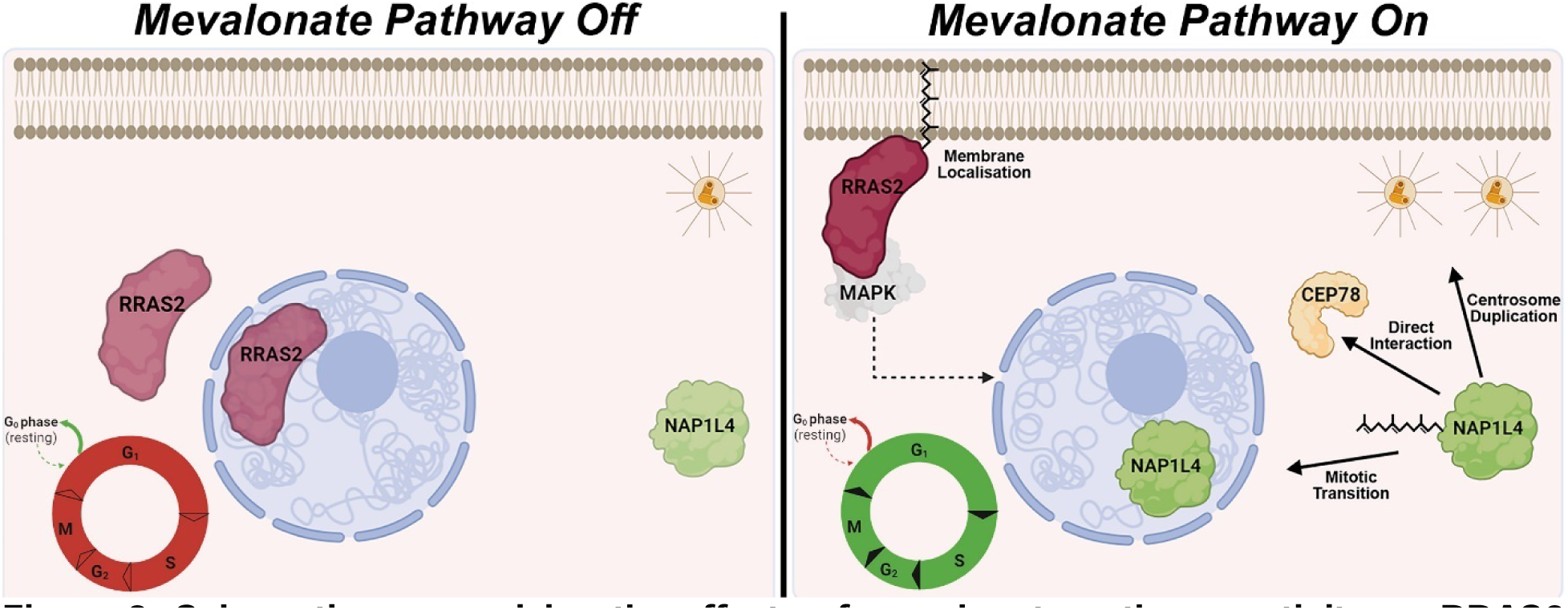
Schematic summarizing the effects of mevalonate pathway activity on RRAS2 and NAP1L4 protein function. When the mevalonate pathway is repressed and CMs are quiescent (left), RRAS2 is found predominantly in the cytoplasm and nucleus whilst NAP1L4 is cytoplasmic. When the mevalonate pathway is active (right), CM cell cycle is enhanced. For RRAS2, prenylation allows localization to the plasma membrane facilitating protein activation. For NAP1L4, prenylation facilitates interaction with CEP78, duplication of centrosomes in CMs and mitotic transition.

This study provides supporting evidence for prenylation as a major pathway for mevalonate pathway-induced proliferation. These findings are consistent with previous data showing that only GGPP could rescue proliferation in statin-treated hPSC-CMs whereas squalene/cholesterol could not [10]. This role for geranylgeranylation was also observed *in vivo* during murine cardiac development with CM-specific knockout of *GGPPS* resulting in reduced CM cell cycle and mitosis [60]. In other cell types, a similar phenomenon has been observed with GGPP rescuing statin-induced inhibition of YAP/TAZ which is required for proliferation in breast cancer cells [61]. Similarly, GGPP supplementation ablated the anti-proliferative effects of a GGPPS inhibitor in leukaemia cells [62]. However, in both studies, FPP addition did not reverse the effects elicited by the inhibitor, whereas in this study we show that FPP rescues hPSC-CM cell cycle. These differences may be cell-type specific potentially due to the farnesylated proteins in CMs differing from these cancer cell types or the dysregulated metabolic and cell cycle state present in cancer cells [63]. From the siRNA experiments performed in this study, it was clear that both FPP and GGPP are acting as substrates for prenylation and this promotes hPSC-CM proliferation.

The role of prenylation in proliferation has been of significant interest, as many oncogenes are reliant on prenylation for activation [64]. Yet the ability to identify and characterize all proteins capable of prenylation at a proteome-wide scale has lagged. Improvement of chemical proteomic techniques has recently enabled “prenylomes” to be generated for endothelial cells [38] and brain-derived cells including neurons, astrocytes and microglia [65]. To determine which specific prenylated proteins may be responsible for CM proliferation, we used two approaches: 1) mass spectrometry analysis of the top proliferation hits in our recent drug screen [10], and 2) alkyne-tagged prenyl probes and mass spectrometry to determine what prenylated proteins are present in hPSC-CMs. This allowed us to identify 40 prenylated proteins enabling the direct identification of prenylated proteins at endogenous abundance at the whole proteome-level in cardiomyocytes. Many of these proteins may play a key role in cardiac proliferation, but we provide direct evidence of the role of NAP1L4 and RRAS2 for hPSC-CM proliferation.

For RRAS2, a role in proliferation has been observed in other non-cancerous cell types including epithelial cells [66], as well as B and T cells [67] and is likely via ERK activation and MAPK signaling in these cell types [54, 68]. Increased RRAS2 expression has also been observed in multiple carcinomas [69] and shown to be required for their proliferation [70]. In a recent paper studying border zone onset after myocardial infarction, RRAS2 was upregulated in a specific border zone CM population [44]. Furthermore, activating RRAS2 mutations can cause Noonan Syndrome, which is associated with congenital heart defects [54] and CM proliferation abnormalities [71]. Our data suggest that prenylation localizes RRAS2 to the cell membrane which is an important component of the signaling function of RAS proteins. Interestingly, activation of ERK signaling was independent of prenylation, yet two gain of function RRAS2 mutants (A70T and Q72L) induced ERK phosphorylation which is typically associated with increased proliferation. Thus, gain of function mutations may alter kinase activation profiles rather than over-activate endogenous ones. The role of RRAS2 in signaling for proliferation in the heart therefore requires further interrogation.

We also interrogated another prenylated protein NAP1L4, revealing a previously unknown role in hPSC-CM proliferation. Knockdown of NAP1L4 using RNA interference led to a significant reduction in proliferating hPSC-CMs, consistent with data in both cervical cancer and melanoma where NAP1L4 silencing leads to cell cycle arrest at G1/S [72, 73]. Using RNA-sequencing we found that NAP1L4 knockdown reduced expression of cell cycle genes. This analysis did not identify any specific signaling pathway as being grossly perturbed, which indicates that NAP1L4 may not be a component of a classical signaling cascade and the reduction of cell cycle genes was most likely via its canonical role as a nuclear chaperone.

Our results reveal a new mechanism of how prenylation regulates hPSC-CM proliferation. Specifically, we show that NAP1L4 prenylation regulates nuclear localization in a cell-cycle specific manner and as such controls centrosomal protein:protein interactions and mitotic transition. Protein pulldown comparing NAP1L4 wild type and prenyl mutant (C383S) interacting proteins uncovered that binding to CEP78 was completely abolished in the mutant. CEP78 is a centriolar protein that has been shown to interact with PLK4 – a kinase required for centriole duplication [74]. If CEP78 expression is inhibited, PLK4 centrosome localization is impaired, thereby reducing centriolar duplication [75]. We have shown that PLK4 inhibition in hPSC-CMs resulted in decreased centrosome numbers, reduced hPSC-CM proliferation and promotion of terminal differentiation [59]. A recent study in human cell lines on separase, which is known to regulate centriole separation at the end of mitosis, also identified NAP1L4 as a centrosomal-interacting partner during mitosis [76]. Here we show that blockade of the mevalonate pathway, knockdown of prenyltransferases and knockdown of NAP1L4 all lead to reduced centrosome number and proliferation in hPSC-CMs. Furthermore, this was specific to NAP1L4 knockdown, as RRAS2 knockdown decreased proliferation, but did not decrease centrosome numbers. These data taken together suggest that prenylated NAP1L4 is directly linked with centrosome dynamics.

NAP1L4 overexpression led to inconsistent CM proliferation in matured hCOs. This likely relates to the changes that occur during cardiac maturation such as reduced CM proliferation [3, 77] and repressed mevalonate pathway expression [27]. Our protein pulldown and live-cell imaging findings highlight the critical role of NAP1L4 prenylation in cell cycle activation and likely explains the lack of proliferative response observed after NAP1L4 overexpression alone. Together our study indicates that prenylation controls multiple cellular processes, and that reactivation of proliferation in a mature cardiomyocyte may require multiple simultaneous processes to be activated. Furthermore, direct activation of the mevalonate pathway via overexpression of master regulators such as SREBF2 are not sufficient to induce CM proliferation. Instead, rewiring of metabolic flux to activate the mevalonate pathway and new techniques to prenylate or mimic prenylation of proteins including NAP1L4 may be required to reactivate proliferation.

In summary, our findings outline prenylation as a branch of the mevalonate pathway responsible for CM proliferation. We identify 40 prenylated proteins in proliferating hPSC-CMs and validate that multiple prenylated proteins play a key role in CM proliferation including (but not limited to) RRAS2 and NAP1L4. Considering the mevalonate pathway is downregulated during cardiac maturation, future work is needed to determine how to effectively reactivate this pathway or artificially “prenylate” critical regulators for cardiac regeneration.

## Methods

### Human Pluripotent Stem Cells

Ethical approval for the use of human embryonic stem cells (hESCs) was obtained from QIMR Berghofer’s Ethics Committee (P2385) and was carried out in accordance with the National Health and Medical Research Council of Australia (NHMRC) regulations. Female HES3 (WiCell) human embryonic stem cells, male CW30382A (designated AA, FujiFilm) and male GM23338 (designated PGP1, Coriell) were maintained in mTeSR Plus (Stem Cell Technologies)/Matrigel (Corning) and passaged using ReLeSR (Stem Cell Technologies). DNA fingerprinting and karyotyping were undertaken for quality control.

### Cardiac Differentiation

Cardiac differentiation was performed as previously described [77–79]. Briefly, hPSCs were seeded onto Matrigel-coated flasks at 2 x 10^4^ cells/cm^2^ and cultured in mTeSR Plus for 3 days and mTeSR1 for 1 day prior to differentiation. To differentiate into cardiac mesoderm, hPSCs were cultured in RPMI B27-medium (RPMI 1640 GlutaMAX, 2% B27 supplement without insulin, 200 µM L-ascorbic acid 2-phosphate sesquimagnesium salt hydrate (Sigma) and 1% Penicillin/Streptomycin (ThermoFisher Scientific) supplemented with 1 µM CHIR99021 (Stem Cell Technologies), 9 ng/mL Activin A, 5 ng/mL BMP4, and 5 ng/mL FGF (RnD Systems). Mesoderm induction required medium changes daily for 3 days. This was followed by cardiac specification using RPMI B27-medium containing 5 µM IWP-4 (Stem Cell Technologies) for another 3 days, and then a further 7 days using 5 µM IWP-4 in RPMI B27+ (RPMI 1640 GlutaMAX, 2% B27 supplement with insulin, 200 µM L-ascorbic acid 2-phosphate sesquimagnesium salt hydrate and 1% Penicillin/Streptomycin) with media changes every 2-3 days. For the final 2 days of differentiation, hPSCs were cultured in RPMI B27+ with no IWP-4. Differentiated cardiac cells were harvested using 0.2% collagenase (Sigma) in 20% Fetal Bovine Serum (FBS) in PBS (with Ca^2+^ and Mg^2+^) at 37°C for 1 hour, followed by exposure to 0.25% Trypsin-EDTA at 37°C for 10 minutes. Cells were filtered through a 100-µm mesh cell strainer (BD Biosciences), centrifuged at 300 x g for 3 minutes, and re-suspended in CTRL medium: α-MEM GlutaMAX, 200 µM L-ascorbic acid 2-phosphate sesquimagnesium salt hydrate (Sigma), 1% Penicillin/Streptomycin and 10% FBS. Previous flow cytometry analysis indicated that differentiated cardiac cells were ∼70% α-actinin^+^/cTNT^+^ cardiomyocytes and ∼30% CD90^+^ stromal cells [79].

### *2D* Cardiac Cell Culture

Day 15 dissociated differentiated cardiac cells were resuspended in CTRL medium and plated at ∼150,000 cells/cm^2^ or 100,000 cells/cm^2^ in a 96-well plate and 6-, 12-, 24-well plates, respectively, that were coated with 0.1% gelatin (Sigma). For all inhibitor and rescue experiments, after 2 days of culture in CTRL medium, cardiac cells were washed with RPMI++ (RPMI 1640, 200 µM L-ascorbic acid 2-phosphate sesquimagnesium salt hydrate, 1% Penicillin/Streptomycin) and cardiac cells were subsequently cultured in RPMI B27+. For 3-day inhibitor experiments, cardiac cells were treated with 10 µM simvastatin (Santa Cruz Biotechnologies), 20 µM FTI-277, 20 µM GGTI-298 or 10 µM TAK-475 (all Sigma). For 1-day rescue experiments, after 2 days of culture in CTRL medium, cardiac cells were treated with 10 µM simvastatin, 20 µM GGTI-298 or 10 µM TAK-475 plus DMSO, 500 µM (±) mevalonic acid 5-phosphate lithium salt hydrate, 20 µM farnesyl pyrophosphate, 20 µM geranylgeranyl phosphate and/or 20 µM cholesterol (all Sigma) 1 day. For 18-hour tipifarnib (Sigma) experiments, after 2 days of culture in CTRL medium, cardiac cells were treated with 1 µM tipifarnib for 18 hours. For prenyl probe experiments, cells were exposed to YnF or YnGG for 16 hours at the concentrations indicated in the figure. For prenyl probe experiments with mevalonate pathway inhibitors, cells were exposed to simvastatin, tipifarnib or GGTI-298 for 1 hour before addition of YnF or YnGG and subsequent culture for 16 hours.

### Identification of Prenylated Proteins in Mass Spectrometry of Proliferation Screen Hits

Our previous proteomics data on proliferation hits was used for this analysis PRIDE: PXD009133 [10]. Targeted data extraction to quantify prenylated peptides was performed manually in the Skyline Environment [80] on Andromeda/MaxQuant search results which included the variable modification of farnesylation (cysteine; C15H24; 204.1878) and geranylgeranylation (cysteine; C20H32; 272.2504) with neutral loss. Data were filtered to 1% FDR at the PSM and peptide level, and only peptides with an Andromeda score >100 and a localization probability >0.75% were included in the analysis.

### Preparation of Cell Lysate Tagged with Prenyl Probes

Cells were put onto ice and washed 2x with ice-cold PBS. Then cells were lysed with lysis buffer (0.2% SDS, 1% NP-40, 150 mM NaCl, 1 mM MgCl_2_, 25 U/mL Benzonase, 50 mM HEPES, pH 7.4) containing complete EDTA-free protease inhibitor cocktail (Roche). Lysate was left on ice for 25 minutes and then shaken at 300 rpm for 10 minutes at room temperature. Lysate was centrifuged at 15,000 x g for 10 minutes at 4°C and supernatant was transferred to a fresh tube and an aliquot taken for quantitation using a BCA assay (Thermo Fisher).

### Click Chemistry General Protocol

Protein amounts were normalized across all samples. For each 100 µL of protein lysate, the click reagent mixture was prepared as follows: 1 µL capture reagent (AzTB or AzRB; 10 mM in DMSO), 2 µL CuSO4 (50 mM in water), 2 µL TCEP (50 mM in water) were added sequentially, mixed and left for 2 minutes at room temperature. Then 1 µL TBTA (10 mM in DMSO) was added. The click reagent mixture was then added to the protein lysate and incubated with shaking at 300 rpm for 1 hour at room temperature. The click chemistry reaction was quenched by addition of EDTA to a final concentration of 5 mM. Two reaction volumes of methanol, 0.5 volume of chloroform and 1 volume of water were added sequentially and mixed by vortexing before leaving on ice for 15 minutes. This was then centrifuged at 15,000 x g at 4°C for 5 minutes. The methanol/water layer followed by the chloroform layer was then aspirated, leaving only the protein pellet. Protein pellet was washed with 500 µL methanol followed by vortexing and sonication for 2 x 5 seconds and left at -80°C overnight. Samples were vortexed and then centrifuged at 15,000 x g at 4°C for 10 minutes. The methanol was then aspirated and the protein pellet was dried on the bench for 5 minutes. The dried pellet was then resuspended in 2% SDS/PBS (1/10th of final volume) followed by vortexing and sonication, before dilution to 0.2% SDS/PBS with PBS.

### In-Gel Fluorescence

Protein (10-16 µg) was combined with 4X Loading Dye and incubated at 95°C for 10 minutes. Denatured protein was resolved by 12% SDS-PAGE gel and run at 100V for 10 minutes followed by 150 V for 1 hour. Gels were incubated in 100 mL fixing solution (40% methanol, 10% acetic acid, water) for 5 minutes at room temperature, then washed 3 x with 100 mL water. Gels were stored in water and protected from light until imaging. Gels were imaged on the Amersham ImageQuant Ai800 imager (GE Healthcare) using the Cy3 and Cy5 channels for the TAMRA fluorophore and protein ladder, respectively. Fluorescence intensity quantification was performed on the Cy3 channel using ImageJ and was normalized to lowest concentration of prenyl probe or probe only.

### Sample Preparation for LCMS-based Analysis of YnF and YnGG Labelled Proteins

Protein lysates were subjected to click chemistry with AzRB capture reagent as described. After precipitation and re-suspension, the protein solution was centrifuged (4000 x g, 10 min, RT) to pellet any particulates. The clarified protein samples were incubated with NeutrAvidin® Agarose resin (50 µL per 1 mg protein, Thermo Scientific) for 2 hours at RT. The beads were pelleted (3,000 x g, 3 min) and supernatant was aspirated. The beads were washed sequentially in 1% SDS in PBS (3 x 0.5 mL), 4M Urea in PBS (2 x 0.5 mL) and 50 mM ammonium bicarbonate (5 x 0.5 mL). For each wash step, the beads were gently vortexed for 1 min followed by pelleting in a microcentrifuge (3,000 x g, 2-3 min). After the final wash the beads were re-suspended in 50 mM ammonium bicarbonate (50 µL). DL-dithiothreitol (5 µL, 100 mM in 50 mM ammonium bicarbonate) was added and the beads incubated at 55°C for 30 minutes in a shaker. The beads were washed with 50 mM ammonium bicarbonate (2 x 0.5 mL) with vortexing and pelleting as before, leaving the beads covered in 50 µL solution after the second wash. Iodoacetamide (5 µL, 100 mM in 50 mM ammonium bicarbonate) was added and the beads incubated at room temperature for 30 minutes in the dark. The beads were washed as before. Sequence grade trypsin (5 µL, 0.2 µg/µL in 50 mM ammonium bicarbonate) was added and the beads incubated at 37°C overnight in a shaker. The beads were pelleted, and the supernatant collected.

### LCMS-based Proteomics of Prenyl Probe Samples

Samples were acidified to pH < 3 and cleaned using tips packed in-house with Empore SDB-RPS material. They were then dried on a speedvac and reconstituted in 0.1% formic acid (FA) for LCMS analysis.

Samples were loaded on to a Thermo Acclaim PepMap 100 trap column (5 mm x 300 µm ID) for 5 min at a flow rate of 10 µl/min with 95% Solvent A (0.1% FA in water) and subsequently separated on a Thermo PepMap100 analytical column (150 mm x 300 µm ID) equipped on a Thermo Ultimate 3000 LC interfaced with Thermo Exactive HF-X mass spectrometer. Peptides were resolved using a linear gradient of 5% solvent B (0.1% FA in 80% acetonitrile) to 40% solvent B over 48 min at a flow rate of 1.5 µL/minutes. This was followed by column washing and equilibration for a total run time of 65 minutes. Mass spectrometry data was acquired in positive ion mode. Precursor spectra (350-1400 m/z) were acquired on the Orbitrap at a resolution of 60,000. The AGC target was set to 3×10^6^ with a maximum ion injection time of 30 ms. Top 20 precursors were selected for fragmentation in each cycle and fragment spectra was acquired in the Orbitrap at a resolution of 15,000 with stepped collision energies of 28, 30 and 32. The AGC target was 1×10^5^, with a maximum ion injection time of 45 ms. The isolation window was set to 1.2 m/z. Precursors with charge states from 2-7 were selected for fragmentation.

Raw LCMS data was searched against the reviewed UniProt human database (20,399 sequences, downloaded April 2021) using Sequest HT on the Thermo Proteome Discoverer software (Version 2.2). Precursor and fragment mass tolerance were set to 20 ppm and 0.1 Da, respectively. A maximum of two missed cleavages were allowed. A strict false discovery rate (FDR) of 1% was used to filter peptide spectrum matches (PSMs) and was calculated using a decoy search. Carbamidomethylation of cysteines was set as a fixed modification, while oxidation of methionine and N-terminal acetylation were set as dynamic modifications. Exploration of ontologies using identified prenylated proteins were performed using Enrichr web interface [40–42] using GO Biological Process 2023 and GO Cellular Component 2023.

### Bioinformatics Analysis to Identify CaaX Motif-Containing Proteins

The NCBI Human RefSeq Protein database (GRCh38) was downloaded and filtered using a custom written script using RStudio (v 3.6.2). Briefly, proteins containing a cysteine “C” residue at the 4^th^ last residue of the protein sequence were filtered. Next, proteins ending in prenylation-specific amino acids [39] were filtered for with the last residue being a “A, V, P, I, L, F, M, S, C, T, or Q” and the second last residue not a “D, E, K, R, Y or W”. Finally, protein isoforms that did not differ in the CaaX motif were removed.

### Modified mRNA Synthesis

Production of in vitro transcription (IVT) template and subsequent modified mRNA (modRNA) synthesis was achieved by combining two previously published protocols [81, 82]. All oligonucleotide constructs were manufactured by Integrated DNA Technologies (Coralville, USA or Singapore). ORFs with UTRs were amplified by PCR (Table S11) and had SacI and SalI restriction enzyme sites introduced. PCR reactions were carried out with Platinum Taq DNA Polymerase High Fidelity (ThermoFisher Scientific) according to the manufacturer’s instructions. Reactions were then purified and concentrated using QIAquick PCR Purification Kit (Qiagen) according to the manufacturer’s instructions. pBluescript II KS (+) and ORF were digested using SacI and SalI (New England Biolabs) restriction enzymes according to the manufacturer’s instructions. Reactions were purified using QIAquick PCR Purification Kit. Ligation of pBluescript II KS (+) and ORF was performed using T4 DNA Ligase (New England Biolabs) according to the manufacturer’s instructions. Ligated products were transformed into Subcloning Efficiency DH5α Competent Cells (ThermoFisher Scientific) according to the manufacturer’s instructions and bacteria were cultured overnight on agar plates with ampicillin (Sigma). Single colonies were picked and screened using PCR to detect presence of insert and cultured overnight in Lysogeny Broth with ampicillin. Plasmid DNA was extracted from bacteria using QIAprep Spin Miniprep Kit (Qiagen) according to manufacturer’s instructions. Plasmid DNA was linearized using AgeI restriction enzyme (New England Biolabs) then Poly A Tail was introduced using PCR (Table S11). modRNA was synthesized using MEGAscript T7 Transcription Kit (ThermoFisher Scientific) with 1.6 µg of DNA template used per 40 µL reaction with a ribonucleoside mix containing 10 mM ARCA (Trilink Biotechnologies), 2.7 mM GTP, 8.1 mM ATP, 8.1 mM CTP, and 2.7 mM N^1^-Methylpseudo-UTP (Trilink Biotechnologies). After incubation at 37°C for 6-8 hours (dependent on construct length), 2 µL Turbo DNase was added and incubated at 37°C for 15 minutes. The modRNA was purified using MEGAclear Kit (ThermoFisher Scientific) according to the manufacturer’s instructions and then treated with Antarctic Phosphatase (New England Biolabs) and incubated at 37°C for 1 hour. ModRNA was again purified using MEGAclear kit and quantitated using NanoDrop. ModRNA molecular weight was verified using denaturing agarose gel electrophoresis.

### Modified mRNA Transfection

Day 15 dissociated differentiated cardiac cells were resuspended in CTRL medium and seeded in 0.1% gelatin-coated plates. After 2 days of culture, cells were washed with DMEM+++ (DMEM without glucose, glutamine and phenol red (ThermoFisher Scientific), 1% GlutaMAX (ThermoFisher Scientific), 200 µM L-ascorbic acid 2-phosphate sesquimagnesium salt hydrate and 1% Penicillin/Streptomycin) and supplemented with 2D maturation medium (DMEM+++, 4% B27 supplement minus insulin, 33 µg/mL aprotinin, 1 mM glucose, 100 µM palmitic acid (Sigma) (conjugated to bovine serum albumin in B27) for 3 days. For transfection, 1 µg or 4 µg of modRNA was mixed with Opti-MEM reduced serum medium plus GlutaMAX (ThermoFisher Scientific) and Lipofectamine RNAiMAX (ThermoFisher Scientific) as per manufacturer’s instructions for a single well of 96-well or 12-well tissue culture plates, respectively. modRNA/lipid complex was added to cells 1:10 with 2D maturation medium and gently rocked to ensure even spreading of transfection complex. For proliferation experiments, cardiac cells were cultured for 48 hours whilst for qPCR and pERK1/2 assay, cardiac cells were cultured for 4 hours. For live cell imaging hPSC-CMs were transfected with GFP-NAP1L4 and GFP-NAP1L4-C383S and cultured for 8 hours before live cell imaging begun. Live imaging was performed in a Leica DM6 microscope and Leica Thunder imager with environmental control at 37°C and 5% CO_2_ with images taken every 2 minutes for 20 hours and analysed using LAS X software.

### qPCR

Total RNA was isolated using the RNeasy Micro Kit (Qiagen) according to the manufacturer’s instructions. Once isolated, NanoDrop 2000 spectrophotometer (ThermoFisher Scientific) was used to quantify and assess purity of RNA. RNA was reverse transcribed using the random primers method in the SuperScript III First-Strand Synthesis System kit (ThermoFisher Scientific) according to the manufacturer’s instructions. The RT-PCR reactions were performed by ABI QuantStudio 5 (Applied Biosystems) using 5 µL of PowerUp SYBR Green Master Mix (Applied Biosystems), 0.5 µL of each oligonucleotide primer, and 1 µL cDNA to a final volume of 10 µL. The 2-ΔΔCt method [83] was used to determine gene expression changes using 18S as a housekeeping gene. Primers for target genes (Table S12) were designed and synthesized by IDT Technologies and were used at 200 nM.

### *2D* Cardiac Cell Immunostaining

Cardiac cells were fixed for 30 min with 1% paraformaldehyde (PFA) (Sigma) at room temperature before being washed twice with PBS. After being washed, cells were subjected to Blocking Buffer (5% FBS and 0.2% Triton X-100 (Sigma) in PBS) for 30 minutes. Primary antibodies α-actinin (1:1000, A7811, Sigma), Ki-67 (1:400, 9129S, Cell Signaling Technology), HA-Tag (1:1000, 3724S, Cell Signaling Technology) and/or pericentrin (1:2000, ab4448, Abcam) diluted in Blocking Buffer were added to cells and incubated for 60 minutes at room temperature. Cells were washed twice with Blocking Buffer before addition of secondary antibodies goat anti-mouse IgG Alexa Fluor 488 (1:400, A-11001), goat anti-rabbit IgG Alexa Fluor 555 (1:400, A-21429) and Hoechst 33324 (1:1000, H3570, all ThermoFisher Scientific) (diluted in Blocking Buffer) for 45 min at room temperature in darkness. Finally, cells were washed twice with Blocking Buffer then left in PBS at 4°C protected from light until imaging.

### *2D* Cardiac Cell Immunostaining Analysis

Imaging was undertaken using ANDOR WD Revolution Spinning Disk confocal microscope with MetaMorph software, Zeiss 780-NLO confocal microscope with Zen Black software, or Leica DM6 microscope and Leica Thunder imager with LAS X software. Images were then analyzed using custom batch processing files written in Matlab R2018a (Mathworks). Cardiomyocytes were distinguished as α-actinin^+^ cells, with nuclei identified as Hoechst 33342^+^. Proliferating cells were identified using Ki-67^+^. The average size of the cardiomyocytes was also calculated. For centrosome analysis, ImageJ was used to manually trace and count cardiomyocyte centrosomes and nuclei under blinded conditions.

### hCO Fabrication

hCO culture inserts were fabricated using SU-8 photolithography and PDMS molding as described previously [77] with the hCO protocol based on [46]. Briefly, acid-solubilized bovine collagen I (Devro) was salt balanced using 10X DMEM (ThermoFisher Scientific) and pH neutralized using 0.1 M NaOH before combining with Matrigel and the cell suspension on ice. Each hCO contained 5 x 10^4^ cardiac cells, a final concentration of 2.6 mg/mL collagen I and 9% Matrigel. A mixture volume of 3.5 µL was pipetted into the hCO culture insert and incubated at 37°C with 5% CO_2_ for 45 minutes in order to gel. After gelling, hCOs were cultured in α-MEM GlutaMAX, 200 µM L-ascorbic acid 2-phosphate sesquimagnesium salt hydrate, 1% Penicillin/Streptomycin, 4% B27 supplement with insulin, 10 ng/mL FGF2 and 10 ng/mL PDGF-BB (R&D Systems) for 2 days. hCOs were subsequently cultured in hCO maturation medium (DMEM+++, 4% B27 supplement minus insulin, 33 µg/mL aprotinin, 1 mM glucose, 100 µM palmitic acid, 10 ng/mL FGF2 and 10 ng/mL PDGF-BB) with medium changes every 2-3 days for 5 days. hCOs were then cultured in weaning medium with medium changes every 2-3 days for 5 days. For mevalonate supplementation experiments, hCOs were treated with either 500 µM (±) mevalonic acid 5-phosphate lithium salt hydrate or vehicle control (water) for 2 days. For NAP1L4 overexpression, hCOs were transduced with 0.625 x 10^10^ AAV6 per hCO encoding NAP1L4 isoform 2 (accession number: NP_001356309.1) or GFP (negative control) under the control of a cardiac troponin T promoter (both from Vector Biolabs) for 4 days.

### Force Analysis of hCO in Heart-Dyno

The pole deflection was used to approximate the force of contraction as per our previous publication [77]. Briefly, time-lapse images were taken with Leica DM6 microscope and Leica Thunder imager with LAS X software for 10 seconds at 50 fps of each hCO under environmentally controlled conditions at 37°C with 5% CO_2_. Custom batch processing files [77] were written in Matlab R2018a (Mathworks) and facilitated the analysis of the contractile properties of the hCOs and the production of time-force graphs [77]. From this, important functional parameters were derived including force, rate and the activation and relaxation times of hCO contraction.

### hCO Immunostaining

hCOs were fixed with 1% paraformaldehyde (Sigma) at room temperature for 1 hour and washed twice with PBS. After fixing, hCOs were subjected to primary antibodies α-actinin, Ki-67 and/or pHH3 (1:200, 06-570, Merck Millipore) diluted in Blocking Buffer and left to oscillate at 45 rpm on a rocker overnight at 4°C. hCOs were then washed twice with Blocking Buffer and oscillated at 4°C for 2 hours. Secondary antibodies goat anti-rabbit IgG Alexa Fluor 555, goat anti-mouse IgG Alexa Fluor 633 (1:400, A-21050, ThermoFisher Scientific) and Hoechst (1:1000) (diluted in Blocking Buffer) were added and left to oscillate overnight at 4°C in darkness. hCOs were then washed twice with Blocking Buffer and oscillated at 4°C for 2 hours. Blocking buffer was removed and hCOs were left in PBS, covered in aluminum foil and stored at 4°C until imaging. After whole tissue imaging, hCOs were mounted onto glass slides (Trajan) using ProLong Glass Antifade Mountant (ThermoFisher Scientific) for high-magnification imaging.

### hCO Immunostaining Analysis

Whole tissue imaging was performed using Leica DM6 microscope and Leica Thunder imager with LAS X software. Custom batch processing files were written in Matlab R2018a (Mathworks) to remove background, calculate image intensity for Ki-67 analysis. For high-magnification imaging of pHH3^+^ mitotic CM, this was performed using a Zeiss 780-NLO confocal microscope with Zen Black software. For analysis, three random field of views per hCO were imaged and manually quantified for mitosis (with sarcomere breakdown) using ImageJ. These were added together to calculate the percentage of CM mitosis in each hCO.

### RNA Interference

Day 15 dissociated differentiated cardiac cells were resuspended in CTRL medium and seeded in 0.1% gelatin-coated plates. After 1 day of culture, cells were transfected for 18 hours with siRNA (ThermoFisher Scientific) according to the manufacturer’s protocol. Briefly, 1 µL gene-specific siRNA oligomer (20 µM) was diluted in 49 µL Opti-MEM reduced serum medium plus GlutaMAX (ThermoFisher Scientific) and mixed with 3 µL of Lipofectamine RNAiMAX (ThermoFisher Scientific) transfection reagent prediluted in 47 µL Opti-MEM reduced serum medium plus GlutaMAX. After incubation for 10 minutes at room temperature, 10 µL of complex was added to a single well of a 96-well plate to a final volume of 100 µL of Opti-MEM reduced serum medium plus GlutaMAX (volumes were upscaled accordingly for larger sized well-plates). After 18 hours, cells were washed with RPMI++ and subsequently cultured in RPMI B27+ for 3 days. For RNA-sequencing experiments, cells were transfected and cultured for 48 hours before harvesting.

### Bulk RNA-sequencing

Total RNA was isolated using the RNeasy Micro Kit (Qiagen) according to the manufacturer’s instructions. Once isolated, RNA was quantified using the Qubit RNA High Sensitivity Assay (Invitrogen). RNA integrity was assessed using the 4200 TapeStation with the RNA ScreenTape kit (Agilent). Samples were normalized to ≤100ng RNA using nuclease free water in a total of 25µl. Poly(A)-enriched RNA libraries were prepared using the Illumina Stranded mRNA Prep, Ligation kit and the IDT for Illumina RNA UD Indexes Set A (UDP0001-UDP0032) (Integrated DNA Technologies). Libraries were quantified using the Quant-iT dsDNA, high sensitivity assay kit (Invitrogen) and library quality was assessed using the QIAxcel DNA High Resolution Kit (1200) (Qiagen). Single-end 100 bp sequencing was conducted on the NovaSeq 6000 using the SP flowcell (Illumina). The knockdown samples (n = 12) achieved a median of 57.18 million reads (range 42.83 to 78.48 million).

Sequence reads were trimmed for adapter sequences using Cutadapt (version 1.9) [84] and aligned using STAR (version 2.5.2a) [85] to the GRCh38 assembly with the gene, transcript, and exon features of Ensembl (release 110) gene model. All read-group aligned BAM files were merged using Samtools merge (version 1.9) for each sample. Quality control metrics were computed using RNA-SeQC (version 1.1.8) [86] and expression was estimated using RSEM (version 1.2.30) [87].

Further RNA-seq analysis was performed in R (version 4.2.0) [88] for protein-coding genes only. The *filterByExpr* function from edgeR (version 3.40.2) [89] was used with default settings for filtering out genes with low counts. Library scaling factors were calculated using edgeR’s calcNormFactors function using the Trimmed Mean of M-values (TMM) method and normalization was performed using edgeR’s cpm function with log set to TRUE to obtain log2 counts-per-million (CPM) for visualization purposes only. The prcomp function from the stats package was used for principal component analysis (PCA), with transposed log2 CPM used as input, and scale and center arguments both set to TRUE. Differential expression analysis was performed using edgeR’s quasilikelihood pipeline [89–91] with filtered raw counts as input. The glmQLFit function from edgeR was used to fit a quasi-likelihood negative binomial generalized log-linear model to the read counts for each gene. The design matrix was defined by model.matrix(∼0 + Group), where Group is a factor with four levels representing the experimental groups HES3 Neg Control siRNA, PGP1 Neg Control siRNA, HES3 NAP1L4 siRNA, and PGP1 NAP1L4 siRNA. We implemented edgeR’s glmTreat function to test for differential expression of a given contrast relative to the default log2-fold-change (logFC) threshold of log2(1.2). Multiple testing correction was performed by applying the Benjamini-Hochberg method on the p-values to control the FDR. Specifically, the function p.adjust from the stats package was used to determine DEGs based on FDR < 5%. The clusterProfiler [92] package (version 4.6.2) was used to perform overrepresentation analysis of Gene Ontology (GO) terms. First, Ensembl IDs for DEGs were converted to Entrez IDs using the bitr function, with the OrgDb argument set to “org.Hs.eg.db”. GO enrichment analysis of sub-ontology biological process was performed using the enrichGO function with the ont argument set to “BP”, and redundant enriched GO terms were removed using the simplify function with default settings. Finally, the dotplot function from enrichplot (version 1.18.4) was used to create dotplots. Additional exploration of cell cycle, cytokinesis and mitotic spindle ontologies using common down-regulated genes from NAP1L4 siRNA experiments were performed using Enrichr web interface [40–42] using GO Biological Process 2023 and GO Cellular Component 2023 and manually filtered for GO terms containing “cell cycle”, “cytokinesis”, or “spindle.” To construct heatmaps, the scale function with center and scale set to TRUE was used to calculate z-scores from log2 CPM, then values for genes of interest were input to ComplexHeatmap (version 2.14.0) [93, 94]. Scatterplots, PCA plots, and mean dispersion plots were created using ggplot2 (version 3.4.2). Venn diagrams were created using the GOVenn function from GOplot (version 1.0.2).

### Affinity Chromatography Mass Spectrometry

Approximately 1 million 2D hPSC-CMs were transfected with either, 10 µg of modRNA GFP-NAP1L4, GFP-NAP1L4-S383 or a vehicle-only control (RNAiMAX) and 2D hPSC-CM were also generated as a negative control. After 8 hours, medium was removed and cells were washed twice with ice-cold PBS. 0.5 mL of lysis buffer [150 mM NaCl, 50 mM Tris-HCl pH 7.5, 1 mM MgCl2, 5% glycerol, 1% IGEPAL CA-630 and protease inhibitor cocktail (Roche)] was added and cells scraped into individual 1.5mL tubes. Genomic material was precleared by centrifugation for 5 min at 1,000 g and supernatant moved to new tubes. 50 µL of prewashed µMACS anti-GFP microbeads (cat: 130-091-288, in lysis buffer) were incubated for 20 minutes with lysed samples. Beads were pelleted by centrifugation at 21,467 g (max speed) for 15 minutes before placing on DynaMag-2 magnet and the supernatant removed. Bound material was washed twice with chilled immunoprecipitation (IP) buffer [150 mM NaCl, 50 mM Tris-HCl pH 7.5, 1 mM MgCl2, 5% glycerol, 0.05% IGEPAL CA-630] followed by three washes with chilled non-detergent IP buffer [150 mM NaCl, 50 mM Tris HCl pH 7.5, 1 mM MgCl2, 5% glycerol] using the centrifuge/magnet separation method. Reduction buffer [2 M Urea, 100 mM Tris HCl pH 7.5, and 10 mM DTT] was added to each sample and incubated for 20 minutes at 37°C, followed by 50 mM Iodoacetoamide in the dark for 30 minutes at 37°C. Samples were diluted with 4 volumes of 100 mM Tris-HCl (pH 7.5), and 350 ng Porcine trypsin (ThermoFisher) added before overnight incubation (17 hours) at 37°C with agitation (1,400 rpm). 20 µL 10% Trifluoroacetic acid was added and sonicated for 15 sec each. Beads were removed by centrifugation, and peptides were dried using GeneVac sample concentrator. Samples were resuspended in equal parts 10% TFA and 5% ACN/0.5% TFA solutions. Self-fabricated SDB-RPS tips were activated and used for desalting, following a standard protocol [95]. Eluted samples were dried and resuspended in 2% ACN/0.3% TFA.

For each sample, a uniform volume (1 µL) was resolved across a 65-minute gradient in data-dependent acquisition mode using a Thermo PepMap100 analytical column equipped on a Thermo Ultimate 3000 LC interfaced with a Thermo Exactive HF-X mass spectrometer. Raw LC-MS data was searched against the reviewed Uniprot human database (20,399 sequences, downloaded April 19, 2021) using Sequest HT on the Thermo Proteome Discoverer software (Version 2.3), with matching between runs enabled. Precursor and fragment mass tolerance were set to 20 ppm and 0.05 Da respectively. A maximum of two missed cleavages were allowed. A strict false discovery rate (FDR) of 1% was used to filter peptide spectrum matches (PSMs) and was calculated using a decoy search. Carbamidomethylation of cysteines was set as a fixed modification, while oxidation of methionine and N-terminal acetylation were set as dynamic modifications. Protein abundance was based on intensity of the parent ions and data was normalized based on total peptide amount. Each test condition (GFP-NAP1L4 or GFP-NAP1L4-C383S) was compared with the negative control protein abundance to determine enriched binding partners of NAP1L4, with a filter cut off of log2FC > 2 in any test condition. An additional filter was used to remove any protein hits detected in only one replicate across the test conditions. Enriched targets were graphically represented using GraphPad Prism (version 9.4.0) as z-score scaled across all samples per protein. Individual proteins were compared across conditions with one-way ANOVA (Tukey multiple corrections, * < 0.05, ** < 0.01, *** < 0.001, *** < 0.0001).

### pERK1/2 Assay

The protocol for AlphaLISA SureFire Ultra pERK1/2 Assay (Perkin Elmer) was followed according to manufacturer’s instructions. Briefly, culture media was removed and plate was incubated at 4°C. 50 µL lysis buffer was added per well and plate was shaken at 1000 rpm for 2 minutes then stored at -20°C until assay was to be performed. After thawing, plate was shaken at 1000 rpm for 1 minute then 10 µL of sample lysate was transferred to a white AlphaPlate-384. Acceptor mix was prepared by combining reaction buffers 1 and 2, 25 x activation buffer and 50 x acceptor beads and 5 µL was added to each well. Plate was sealed and wrapped in foil, shaken at 1000 rpm for 2 minutes, centrifuged at 400 rpm for 20 s then incubated at room temperature for 1 hour. Donor mix was prepared by combining dilution buffer and 50 x donor beads and 5 µL was added to each well. Plate was sealed and wrapped in foil, shaken at 1000 rpm for 2 minutes, centrifuged at 400 rpm for 20 s then incubated at room temperature for 1 hour. Plate was then read at 615 nm using standard AlphaLISA settings on a Biotek Synergy Neo2 (Agilent) plate reader.

### Statistics

All 2D and hCO experiments were performed with multiple replicates per condition in multiple experiments to ensure reproducibility. For non-automated quantification experiments such as centrosome quantification, personnel performing the analyses were blinded to the treatments. For experiments where multiple groups were analyzed, all groups were present in each experiment including controls to ensure that results were not an artefact of comparing conditions over different experiments. Data are presented as mean ± SD unless otherwise noted and statistics were analyzed using Excel (Microsoft) or GraphPAD Prism 8 (Graphpad Software Inc.). Sample numbers, experimental repeats, statistical analyses and p values are reported in each figure legend. Data is presented as n = replicates, from x number of independent experiments.

## Data Availability

The datasets generated and/or analyzed during the current study are available from the corresponding author upon reasonable request and will be publically available following journal publication.

## Author Contributions

C.A.P.B., J.D.R., H.R.R., L.A.C.D., R.Z., E.M., L.T.R., R.L.F., S.R.F., B.L.P., R.J.M., and J.E.H. performed experiments. C.A.P.B., J.D.R., H.R.R., H.C.S., L.A.C.D., R.Z., S.H., L.S., S.M.H., R.L.J., S.R.F., D.C.H.N., B.L.P., R.J.M., and J.E.H. analyzed data. C.A.P.B., E.T., E.R.P., R.J.M., and J.E.H. conceptualized the project. C.A.P.B. and J.E.H. wrote the manuscript. All other authors had input into the manuscript and the final approval.

## Acknowledgements

The Heart-Dyno molds were fabricated using the NCRIS enabled Australian National Fabrication Facility – Queensland, New South Wales and South Australian (Government of South Australia supported) Nodes. We especially thank Dr Simon Doe and Dr Mark Cherrill. We thank Tam Nguyen and Nigel Waterhouse (QIMR Berghofer) for microscopy; Mark Hodson for proteomics support; and Scott Wood and Ross Koufariotis (QIMR Berghofer) for bioinformatic assistance. J.E.H. was supported by funding from a Snow Medical Fellowship. E.R.P. is supported by an Investigator Grant from the National Health and Medical Research Council of Australia (GNT2008376). The Novo Nordisk Foundation Center for Stem Cell Medicine (E.R.P. and R.J.M.) is supported by Novo Nordisk Foundation grants (NNF21CC0073729).

## Supplemental Figure Legends

**Figure S1:**
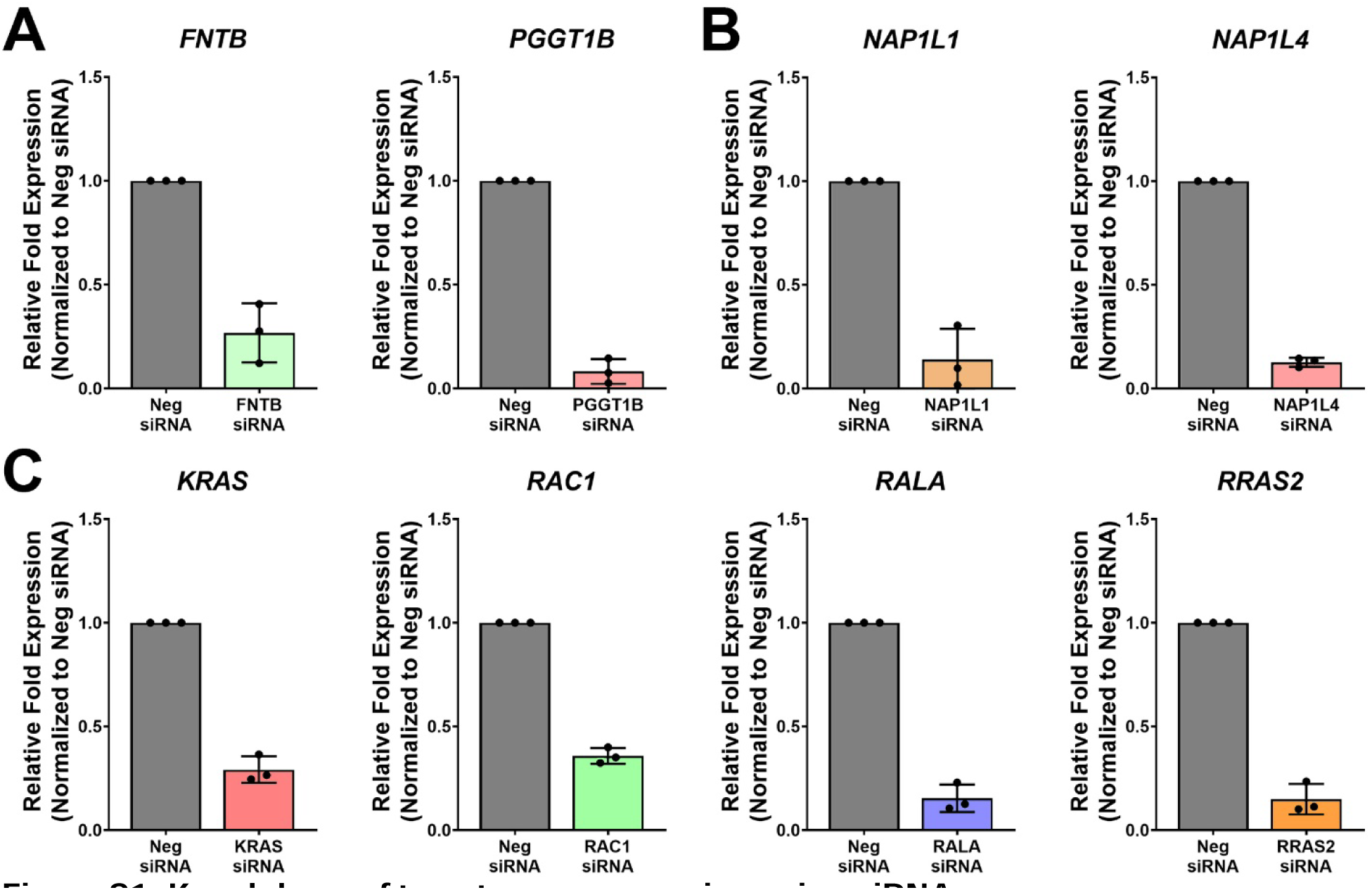
Knockdown of target gene expression using siRNA. Immature 2D hPSC-CMs were transfected with siRNA and cultured for 4 days. The 2-ΔΔCt method was used to determine gene expression changes using 18S as a housekeeping gene with normalization performed relative to the Neg siRNA. n = 3 experiments. Graphs show relative fold expression of target gene for (A) *FNTB* and *PGGT1B*, (B) *NAP1L1* and *NAP1L4*, and (C) *KRAS, RAC1, RALA* and *RRAS2*.

**Figure S2:**
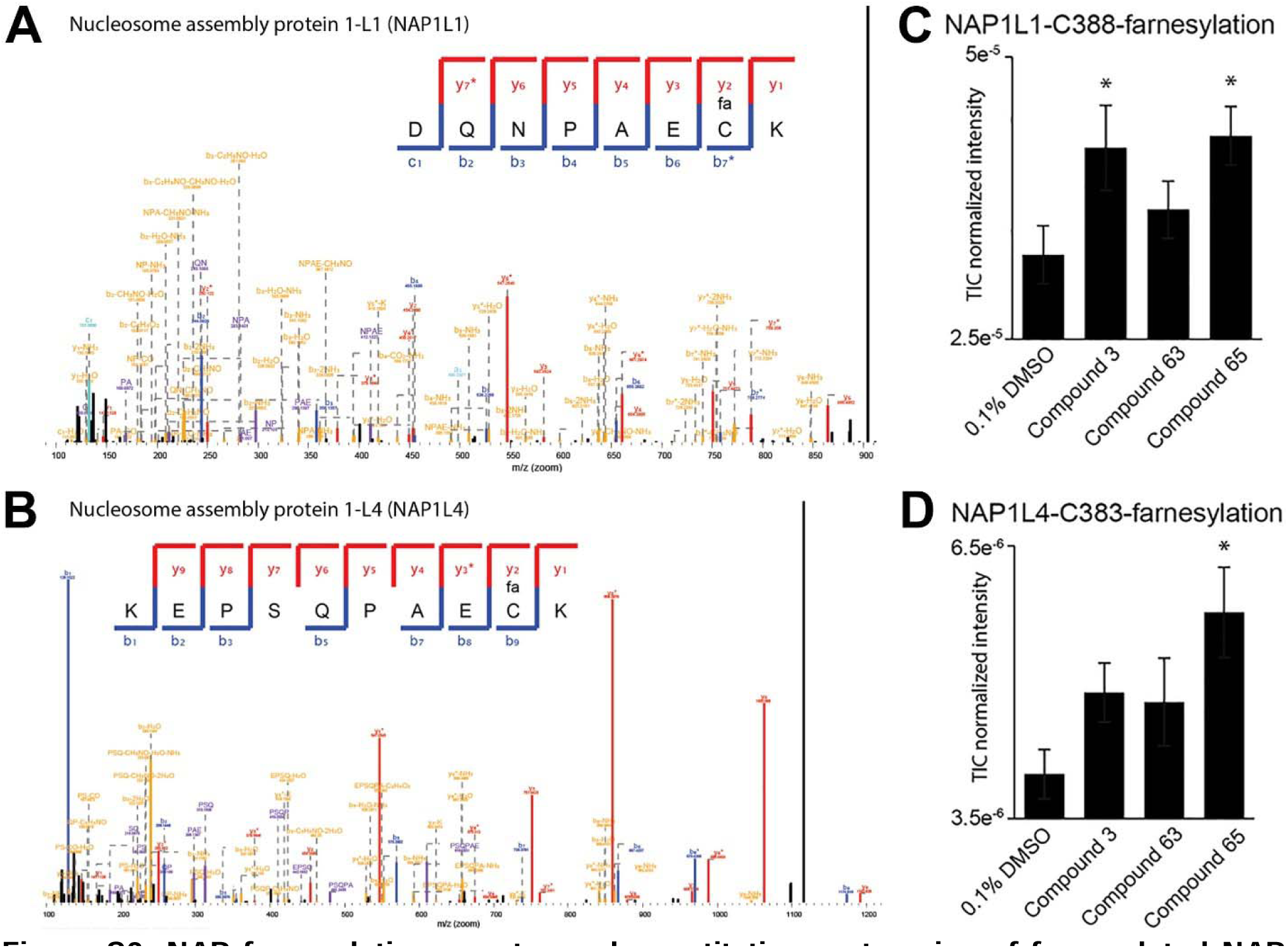
NAP farnesylation spectra and quantitative proteomics of farnesylated NAP proteins after pro-proliferative treatment. (A) Peptide fragmentation map of hCO-derived farnesylated (‘fa” on cysteine residue) NAP1L1 describing precise location and comprehensive coverage of the modification site. (B) Peptide fragmentation map of hCO-derived farnesylated (‘fa” on cysteine residue) NAP1L4 describing precise location and comprehensive coverage of the modification site. (C) Abundance of farnesylated NAP1L1 increased upon pro-proliferative stimulus in hCOs. n = 6 hCOs. TIC = total ion current (D) Abundance of farnesylated NAP1L4 increased upon pro-proliferative stimulus in hCOs. n = 6 hCOs. TIC = total ion current

**Figure S3:**
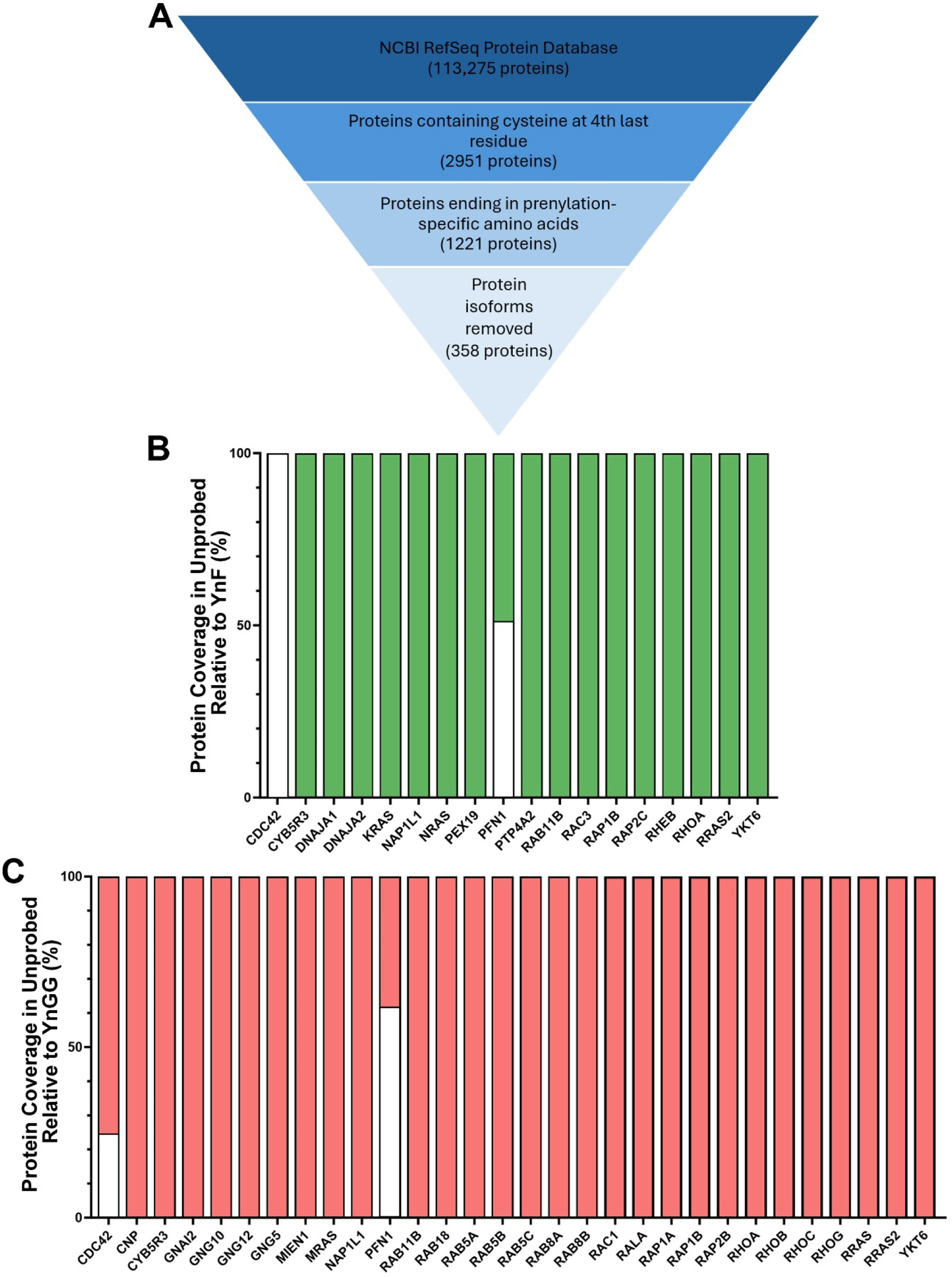
Bioinformatics pipeline for identification of potentially prenylated proteins and protein coverage proportion in unprobed samples relative to probe-treated samples. (A) Bioinformatics pipeline showing the filtering of proteins ultimately identifying 358 potentially prenylated proteins. (B) Proportion of protein coverage of identified farnesylated proteins in unprobed sample (white) relative to YnF sample (green). (C) Proportion of protein coverage of identified geranylgeranylated proteins in unprobed sample (white) relative to YnGG sample (red).

**Figure S4:**
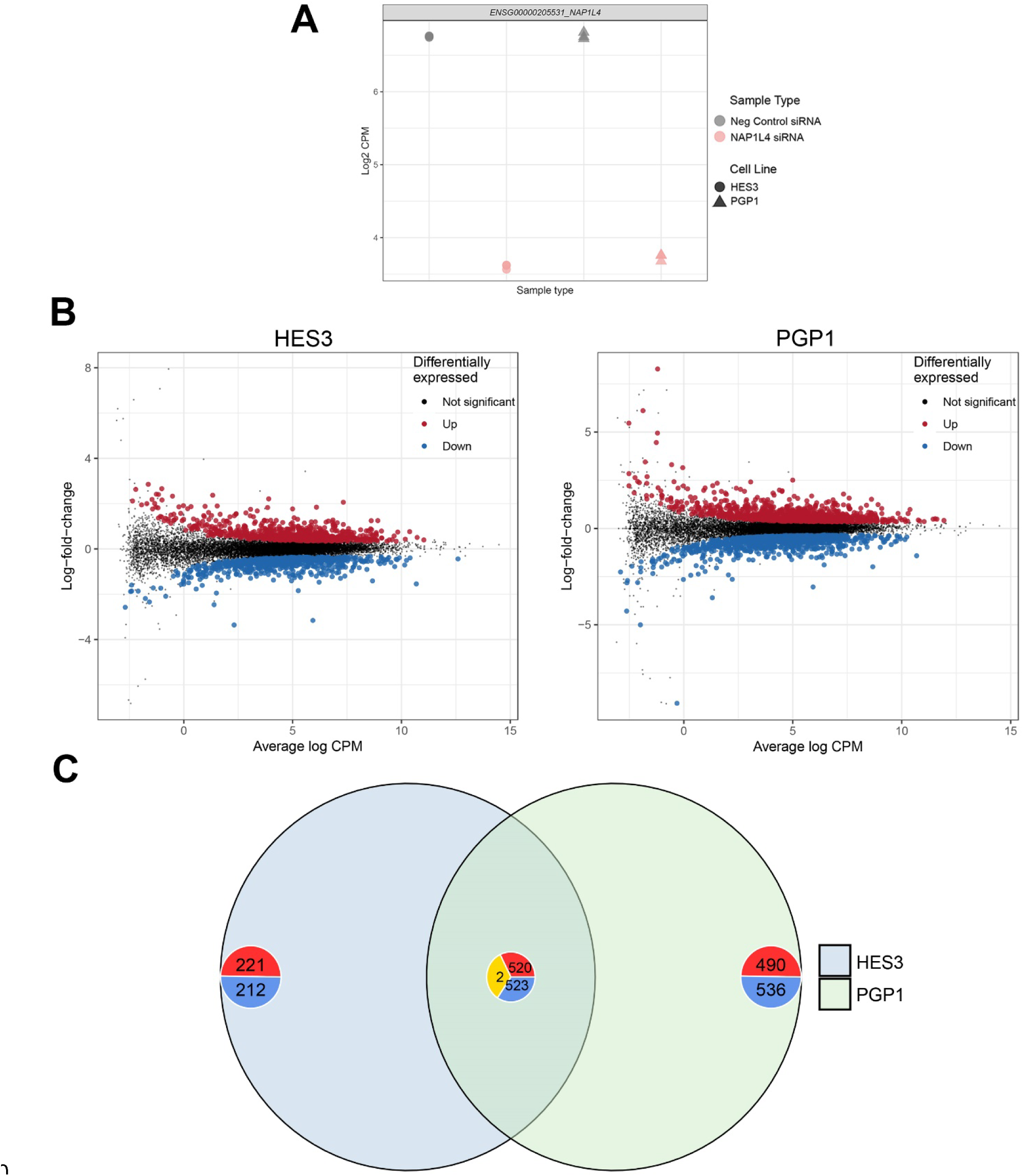
NAP1L4 siRNA treated 2D hPSC-CMs RNA-sequencing plots. (A) Normalized expression (log2 counts per million, CPM) of NAP1L4 siRNA was 47% and 45% reduced after siRNA transfection compared to Neg Control siRNA in HES3 and PGP1 cell lines, respectively. (B) Mean difference plots for differential expression analysis between NAP1L4 siRNA and Neg Control siRNA in HES3 (left) and PGP1 (right). Log2-fold-change vs average log2 CPM. (C) Venn diagram showing comparison of differentially expressed genes between NAP1L4 siRNA and Neg Control siRNA in HES3 and PGP1 cells. Number of up-(red), down-(blue) and oppositely-regulated (yellow) genes displayed.

**Figure S5:**
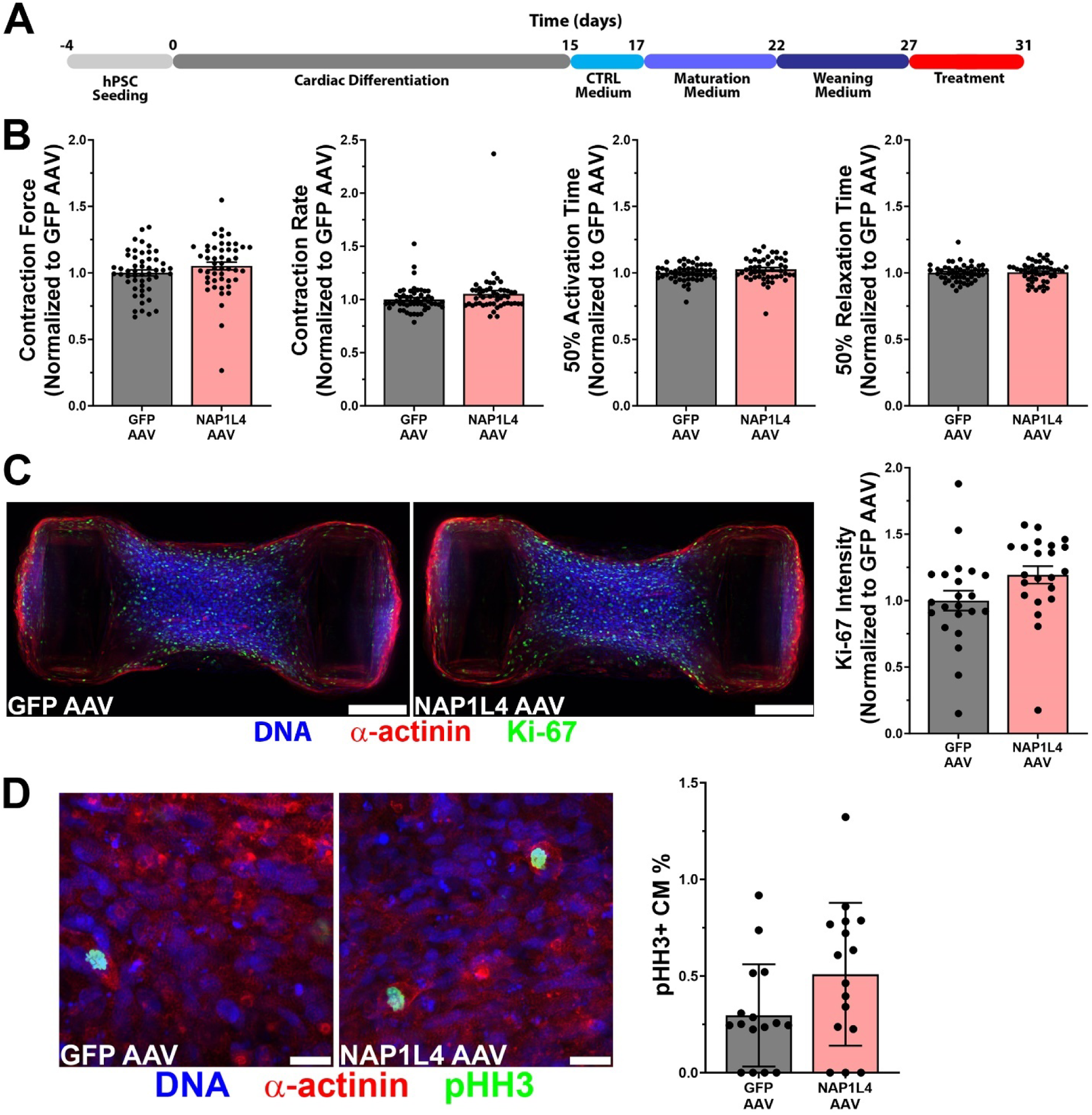
NAP1L4 overexpression has no effect on contractile function or proliferation in mature hCOs. (A) Experimental timeline. Mature hCOs were infected with AAV6-NAP1L4 for 4 days. (B) AAV6-NAP1L4 had no effect on any contractile parameter (force, rate or contractile kinetics). n = 49-51 hCOs from 4 experiments. (C) AAV6-NAP1L4 had no significant effects on Ki-67 intensity. n = 22 hCOs from 4 experiments. Representative images of immunostained hCOs used for whole tissue intensity analysis for Ki-67. Scale bar = 200 µm. (D) AAV6-NAP1L4 did not alter levels of CM mitosis (pHH3+ CMs). n = 16 hCOs from 3 experiments. Representative images of pHH3-stained hCOs to quantify CM mitosis. Scale bar = 20 µm. Data are presented as mean ± SEM (B,C) or ± SD (D).

**Figure S6:**
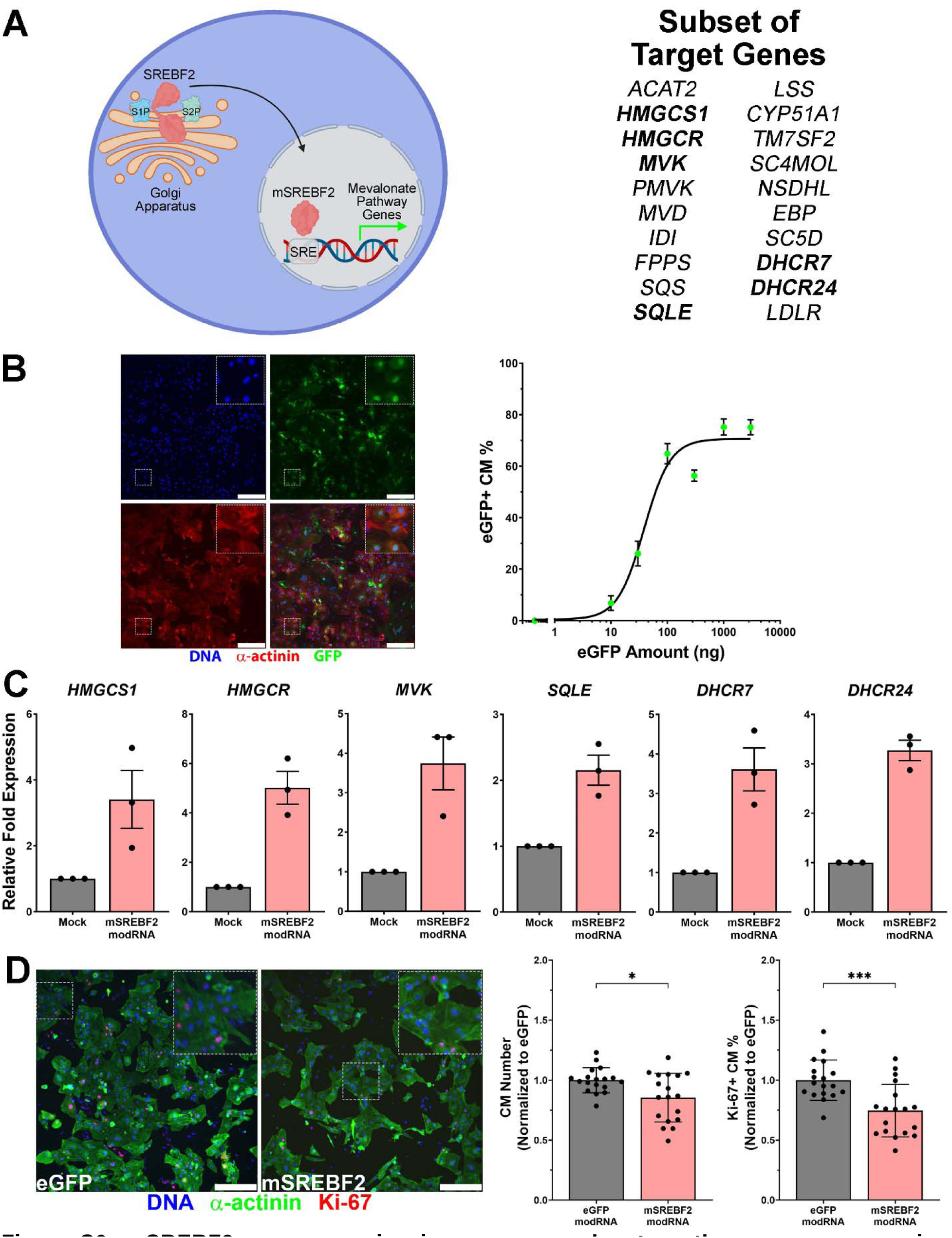
mSREBF2 overexpression increases mevalonate pathway gene expression but reduces CM proliferation in 2D hPSC-CMs. (A) Schematic detailing the mechanism of SREBF2 processing. Inactive SREBF2 resides at the Golgi whereby it is cleaved by site-1(S1P) and site-2 (S2P) proteases allowing the transcriptionally active fragment, mSREBF2, to be translocated to the nucleus. Once in the nucleus, mSREBF2 binds to sterol regulatory element (SRE) sequences in the promoter of target genes and allows transcription. Target genes analyzed in qPCR are indicated in bold text. (B) Representative image of 2D hPSC-CMs transfected with 1000 ng eGFP modRNA for 3 days. Scale bar = 250 µm. EC50 = ∼38.5 ng calculated using the Hill equation. Even though EC50 is ∼38.5 ng, we decided to use 1000 ng modRNA per single well of a 96-well plate as it achieved ∼75% eGFP+ CM nuclei. (C) mSREBF2 modRNA treatment of hPSC-CMs resulted in elevated expression of mevalonate pathway target genes. n = 3 experiments. (D) Representative images of stained mSREBF2 modRNA transfected 2D hPSC-CMs cultured in maturation medium. Scale bar = 250 µm. Treatment of 2D hPSC-CMs cultured in maturation medium resulted in a significant reduction in hPSC-CM nuclei number and cell cycle activity. n = 18 from 3 experiments. Data are presented as mean ± SD. *, *** denotes p < 0.05, p < 0.001, respectively, compared to eGFP modRNA using student’s t-test.

**Figure S7:**
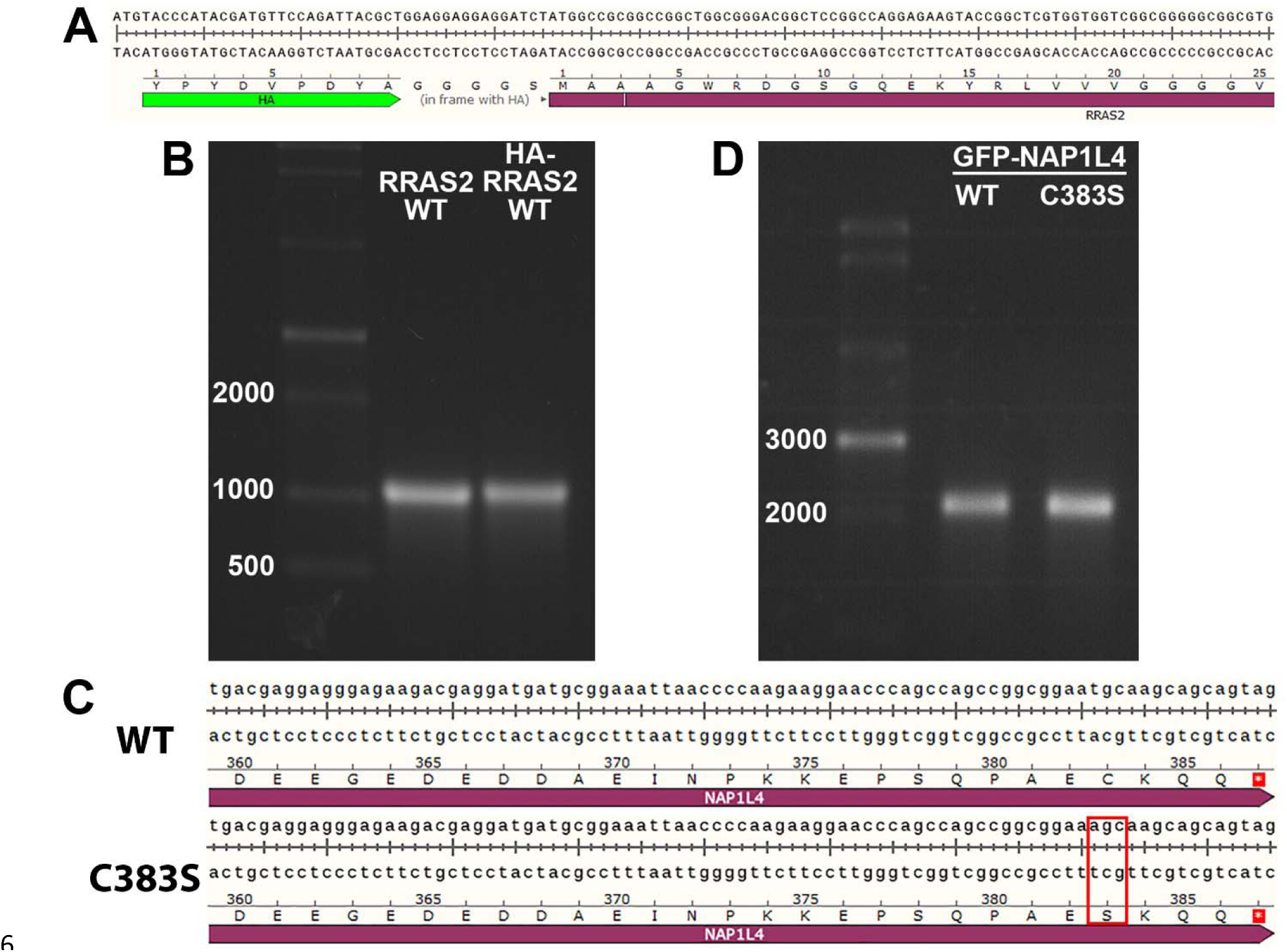
Characterization of tagged NAP1L4 and RRAS2 modRNA. (A) Partial DNA sequence of HA-tagged RRAS2 construct used for modRNA synthesis illustrating insertion of HA tag at the amino-terminus followed by a glycine-serine linker and then the start of the RRAS2 coding sequence. (B) Denaturing agarose gel showing that RRAS2 and HA-RRAS2 modRNA transcript are ∼1000 bp. (C) Partial DNA sequence of GFP-NAP1L4 construct used for modRNA synthesis with red box indicating the cysteine to serine mutation for the prenyl-null mutant at C383 (GFP-NAP1L4-C383S). (D) Denaturing agarose gel showing that GFP-NAP1L4 modRNA transcript is ∼2200 bp.

**Figure S8:**
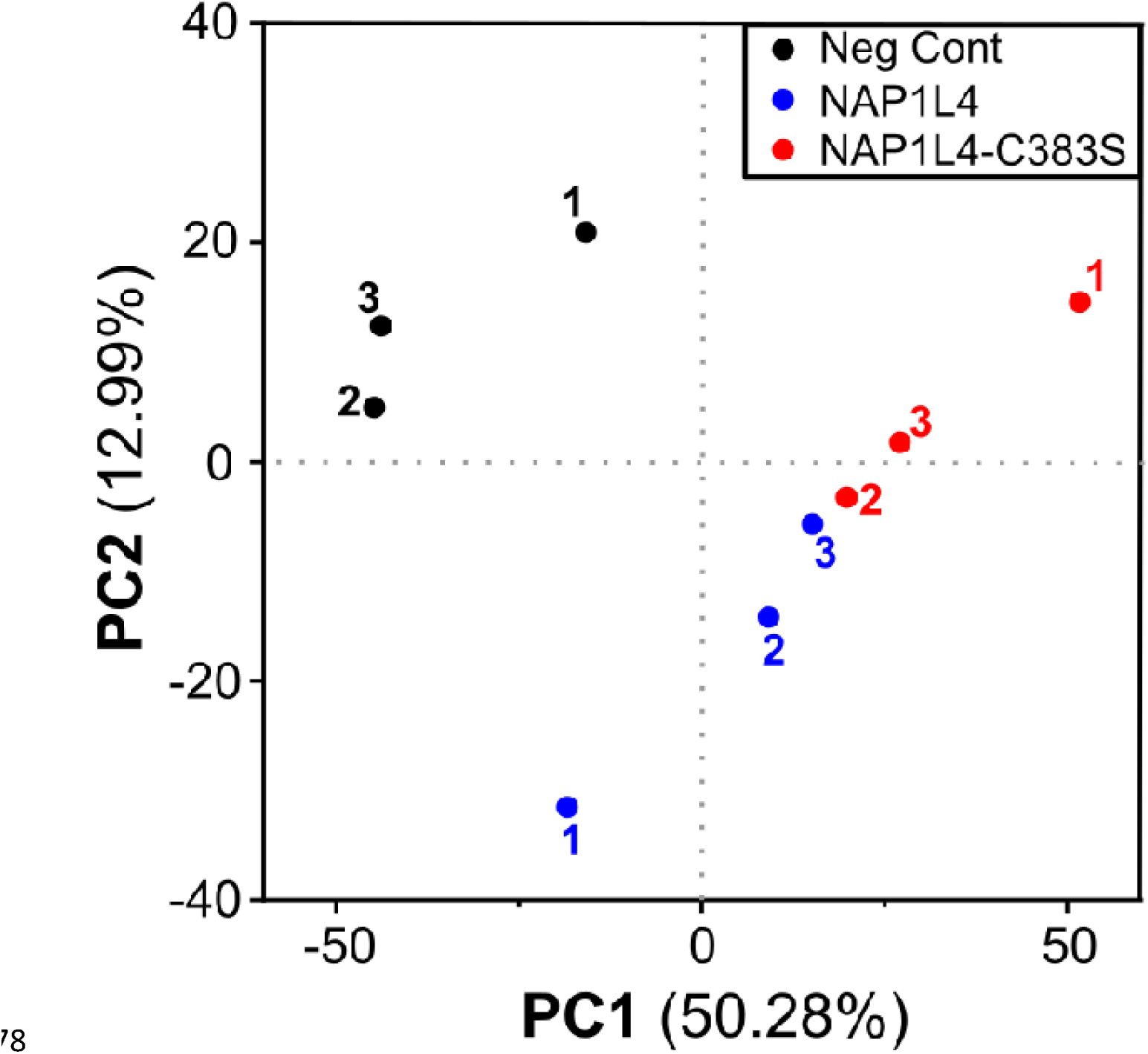
PCA plot of GFP-NAP1L4 modRNA treated 2D hPSC-CMs. PCA of protein pulldown from 2D hPSC-CMs cultured in maturation media transfected with RNAiMAX only (Neg Control), GFP-NAP1L4 or GFP-NAP1L4-C383S modRNA for 8 hours. n = 3 from 1 experiment.

**Figure S9:**
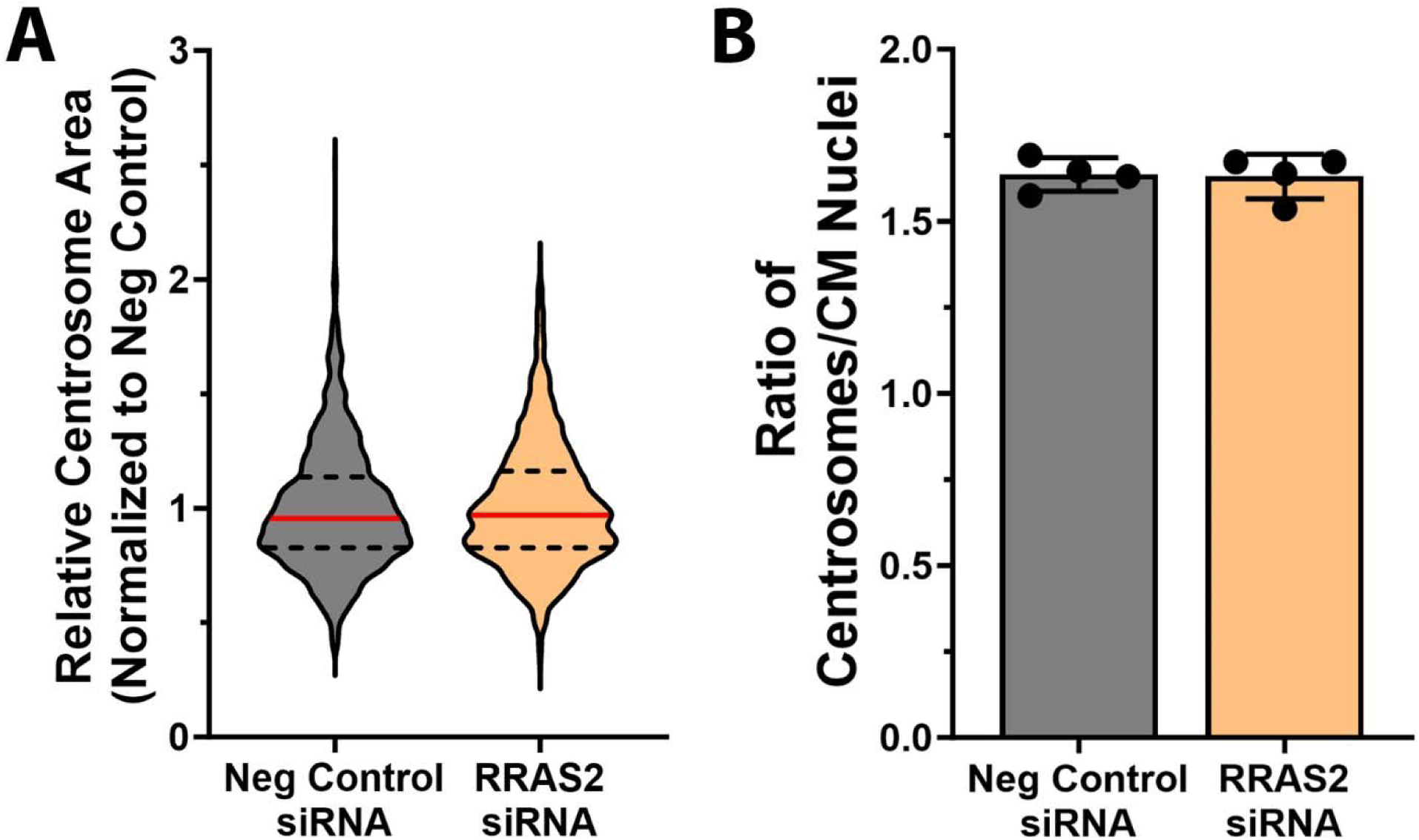
RRAS2 knockdown has no effect on hPSC-CM centrosome size or number. (A) Silencing of RRAS2 did not alter centrosome area in hPSC-CMs. n = 1000-1181 centrosomes from 2 experiments. (B) RRAS2 knockdown had no effect on number of centrosomes per CM nuclei. n = 4 biological replicates over 2 experiments (from 1000-1181 centrosomes total and 609-726 CM nuclei total). Data are presented as median with interquartile range (A) or mean ± SEM (B).

## Supplemental Tables

**Supplementary Table 1: Proteins identified by mass spectrometry in YnF-treated hPSC-CMs.**

**Supplementary Table 2: Proteins identified by mass spectrometry in YnGG-treated hPSC-CMs.**

**Supplementary Table 3: List of 358 potentially prenylated proteins based on presence of carboxyl-terminal CaaX motif.**

As both CDC42 and KRAS have two unique CaaX motifs capable of being prenylated, there are 360 proteins in this list.

**Supplementary Table 4: Enrichr results for gene ontology terms from sub-ontology molecular function for prenylated proteins identified by mass spectrometry.**

**Supplementary Table 5: Enrichr results for gene ontology terms from sub-ontology biological process for prenylated proteins identified by mass spectrometry.**

**Supplementary Table 6: Differential expression analysis results for NAP1L4 siRNA compared to negative control siRNA in 2D hPSC-CMs from HES3 cell line with log2FC (1.2).**

**Supplementary Table 7: Differential expression analysis results for NAP1L4 siRNA compared to negative control siRNA in 2D hPSC-CMs from PGP1 cell line with minimum log2FC (1.2).**

**Supplementary Table 8: Common differentially expressed genes (FDR < 0.05) across PGP1 and HES3 cell lines for respective comparisons between NAP1L4 siRNA and negative control siRNA.**

**Supplementary Table 9: Enriched gene ontology terms from sub-ontology biological process for common down-regulated differentially expressed genes across PGP1 and HES3 cell lines from NAP1L4 siRNA compared to negative control siRNA.**

**Supplementary Table 10: Enrichr results for common down-regulated cytokinesis or mitotic spindle genes after NAP1L4 siRNA.**

**Supplementary Table 11: Primers used for modified mRNA synthesis.**

**Supplementary Table 12: Human primer sequences for qPCR.**

